# Epigenomic and transcriptomic germ-free ageing atlas reveals sterile inflammation as an intrinsic ageing feature

**DOI:** 10.1101/2025.11.19.689100

**Authors:** Ward Nijen Twilhaar, Jen-Chien Chang, Tommy W. Terooatea, Haruka Yabukami, Sachi Kato, Nicola Hetherington, Naoko Satoh-Takayama, Alireza Ahmadi, Tsung-Han Hsieh, Prashanti Jeyamohan, Tomoko Kageyama, Yan Jun Lan, Nanako Kadono-Maekubo, Shiori Maeda, Yurina Miyajima, Miho Mochizuki, Yasutaka Motomura, Yumiko Nakanishi, Nils Rother, Umi Tahara, Natsuki Takeno, Ko Tsutsui, Marina Vilaseca Barcelo, Haitao Wang, Masayuki Amagai, Hironobu Fujiwara, Chung Chau Hon, Takeshi Matsui, Kazuyo Moro, Sussan Nourshargh, Hiroshi Ohno, Aki Minoda

## Abstract

Inflammageing is a hallmark of ageing. Commensal microbiota plays crucial roles in maintaining tissue homeostasis, yet its impact on cellular ageing and inflammageing remains poorly understood. Here we present a comprehensive single-cell epigenomic and transcriptomic atlas of tissues from mice aged under specific pathogen-free (SPF) or germ-free (GF) conditions. Microbiota conferred beneficial effects in young mice but accelerated various ageing features in old, such as age-related AP-1 pathway upregulation, senescence and transcriptomic alterations, likely due to age-associated dysbiosis. Strikingly, inflammatory signatures persisted across cell types in aged GF mouse tissues, establishing sterile inflammation as an intrinsic feature of ageing. Age-associated B cells expanded equally under GF and SPF conditions, raising the possibility that they function as intrinsic, microbiota-independent drivers of inflammageing and potential therapeutic targets. The atlas provides a resource for distinguishing intrinsic ageing features from those modulated by the microbiota, illuminating mechanisms of cellular ageing and potential anti-ageing interventions.

**Highlights:** Sterile inflammation and age-associated B cell expansion are prominent intrinsic ageing features

The upregulation of age-associated AP-1 pathways and senescence of alveolar macrophages are attenuated in germ-free condition

Germ-free condition induces premature ageing-like features in young mouse tissues but delays ageing features in old mice across multiple cell types

## Introduction

Persistent low-grade inflammation with ageing, termed inflammageing, is a hallmark of ageing and is considered a major driver of numerous age-associated diseases [1], [2], [3]. Elucidating and targeting the underlying causes of inflammageing is crucial for promoting healthier longevity. Ageing as a biological phenomenon arises from the interplay between intrinsic cellular processes and extrinsic influences. The relative contribution of intrinsic cellular ageing and extrinsic factors to organismal ageing is currently unclear. Extrinsic factors, such as high-fat diets and persistent infections that alter the tissue microenvironment can accelerate the ageing process [4], [5], [6]. One of the understudied extrinsic factors in the context of ageing is the commensal microbiota, which refers to non-harming microbes such as bacteria, viruses, and fungi, that live on the surfaces of human bodies such as the skin and mucosa. Although Louis Pasteur’s speculation in 1885 that microbes would be essential for organ functions [7] proved incorrect with the development of GF animals, microbiota is still recognised as integral to tissue development and homeostasis, both locally in the mucosa and systemically [8], [9], [10], [11]. On the other hand, age-related imbalances in microbial communities termed dysbiosis has been implicated in inflammageing [12], [13], and transfer of aged microbiota from old specific pathogen-free (SPF) mice to young GF mice under sterile condition promotes inflammageing [14].

Cellular senescence is a tumour-suppressive mechanism characterised by irreversible cell cycle arrest [15] that also increases with old age, and removal of senescent cells (known as senolytics) has been shown to have rejuvenation effects [16], [17]. Senescence is induced by various cell-autonomous states such as high genome instability, mitochondrial dysfunction, and oncogene activations [18], [19]. Key characteristics of senescent cells are their pro-inflammatory state and senescence-associated secretory phenotype (SASP), where production of pro-inflammatory chemokines and cytokines can transform neighbouring cells into senescence resulting in amplification of pro-inflammatory state [20]. Recent evidence suggests dysbiosis may have a causal role in cellular senescence during the course of ageing [21], [22], however, such effect in remote tissues at cellular resolution is still lacking.

In addition to the established hallmarks of ageing [1], recent single-cell transcriptomic profiling has revealed shifts in immune cell composition [23], [24], [25] including GZMK^+^ CD8^+^ T cells in mice and humans [26], cytotoxic CD4^+^ T cells in supercentenarians [27], and innate-like age-associated B cells (ABCs) [28], [29]. Notably, heterochronic parabiosis experiments demonstrate that young blood rejuvenates aged tissues while old blood accelerates ageing in young mice [30], indicating that circulating factors including cells can modulate cellular ageing states. Single-cell epigenomic studies have revealed age-associated remodeling of the chromatin landscape across cell types reinforcing the model that progressive loss of constitutive heterochromatin drives ageing [31], [32], [33], [34], [35]. Other age-related changes include loss of tissue and cellular identity [36], and upregulation of AP-1 pathways and innate immune genes [37].

Here, we present the Mukin ("germ-free" in Japanese) Mouse Ageing Atlas, a comprehensive single-cell epigenomic and transcriptomic resource designed to distinguish intrinsic cellular ageing processes from those modulated by the microbiota. We show that the senescence of alveolar macrophages is significantly accelerated under SPF conditions. These senescent cells adopt a highly inflammatory state, upregulating NF-κB pathway genes, and exhibit metabolic shifts that may contribute to the changing lung microenvironment. Further highlighting the microbiota’s influence, we find the age-associated upregulation of pro-inflammatory AP-1 pathway is significantly attenuated under germ-free conditions, linking microbial presence to this key ageing signature. Importantly, effects of microbiota are context-dependent; we observed that germ-free conditions induced premature ageing-like transcriptomic changes in young, and delayed age-related transcriptomic changes in old mice. Conversely, we identified elevated expression of inflammatory genes across cell types and expansion of ABCs as prominent intrinsic features of ageing that proceeded at the same magnitude in both sterile and SPF conditions across tissues. Together, these findings establish sterile inflammation as a core component of biological ageing and demonstrate how extrinsic factors can selectively exacerbate intrinsic trajectories, providing a valuable resource to disentangle the complex drivers of ageing and ageing-related diseases.

### Tissue ageing features attributable to intrinsic ageing and microbiota

Mice of two ages, young (2 months) and old (19 months), were reared in two conditions: specific pathogen-free (SPF) and GF, which we will refer to as *natural* and *sterile* ageing, respectively (Figure 1A). From these four groups of mice, we generated an ageing atlas of single-cell gene expression (scRNA-seq) and single-cell chromatin accessibility (scATAC-seq). This comprehensive dataset comprised of 180,296 single-cell transcriptomes and 289,489 single-cell chromatin profiles derived from ten whole tissues: bone marrow, peripheral blood mononuclear cells (PBMC), spleen, lung, mesentery, small intestine, large intestine, back skin (epidermis and dermis), and ear skin (epidermis) (Figure 1A-B). These ages are widely used in ageing research and capture key physiological transitions, namely early adulthood and the onset of age-related functional declines, and thus allow us to highlight microbiota-dependent (extrinsic) and -independent (intrinsic) features of ageing across tissues. Dysbiosis that is known to occur with ageing was confirmed in old SPF mice by microbiome analysis of faecal samples, including the decline in diversity and changes in tissue metabolites known to be synthesised in microbiota-dependent manner (Figure S1A-C, Supplementary table 1-3). For example, short chain fatty acids (SCFAs) such as butyrate and propionate that are produced by the gut microbiota by fermentation of dietary fibers were drastically reduced in GF mice, which increased with old age in SPF mice (Figure S1C). Such increases correlate with microbial composition (Figure S1D), although further studies are needed to identify the major sources of such SCFA changes. We found previously unreported declines in nicotinic acid (NA) in old SPF, a metabolite produced in a microbiota-dependent manner [38] (Figure S1E). NA is a precursor of nicotinamide adenine dinucleotide (NAD), which is critical for cellular energy (ATP) production and as co-factors for NAD-dependent enzymes, including signalling molecules, epigenetic and DNA repair enzymes [38]. Such reduced NA level is consistent and correlates with the known decline of cellular NAD levels with age [39]. Although not in a microbiota-dependent manner, another metabolite we detected to decline with age in the intestines as well as remote tissues (lung and skin) was inosine, which is a known anti-inflammatory metabolite (Figure S1E) [40], [41]. We performed clustering and annotation on the full scRNA-seq atlas (Figure S1F-G, Supplementary table 4) and further subclustered for compartments that were initially less separated (Figure S1H-M). Fibroblasts, endothelial cells, and intestinal epithelial cells were further separated by tissue due to tissue-specific differences that arise with their tissue of origin. scATAC-seq data were annotated by transferring labels from our scRNA-seq atlas and by manual curation (Figure S1N-O, Supplementary table 5, Methods). Notably, ∼70% of the ATAC peaks were identified additionally to the ENCODE database (Figure S1P). Disrupted tissue homeostasis is a hallmark of ageing and often appears as shifts in cell type composition, particularly among immune cells. Remarkably, we observed that most cell types exhibited similar compositional changes during both natural and sterile ageing, albeit with variations in magnitude (Figure 1C and S1Q). We defined such conserved features across natural and sterile ageing as “intrinsic” to cellular ageing features, i.e., those that occur irrespective of environment. Notably, ABCs, also known as atypical B cells) showed the largest and most systemic intrinsic ageing-associated abundance changes (Figure 1C). Elevations of plasma cells were particularly prominent in the spleen, and we observed similar but more modest trends for memory B cells and CD8 effector T cells. The relative abundance of naive CD4 effector T cells increased in most tissues during both natural and sterile ageing, except in PBMCs where they decreased during sterile ageing (Figure 1C). Opposite trends in abundance changes between natural and sterile ageing were observed most frequently in the bone marrow, including CD4 and CD8 native T cells, as well as ILCs.

**Figure 1.**
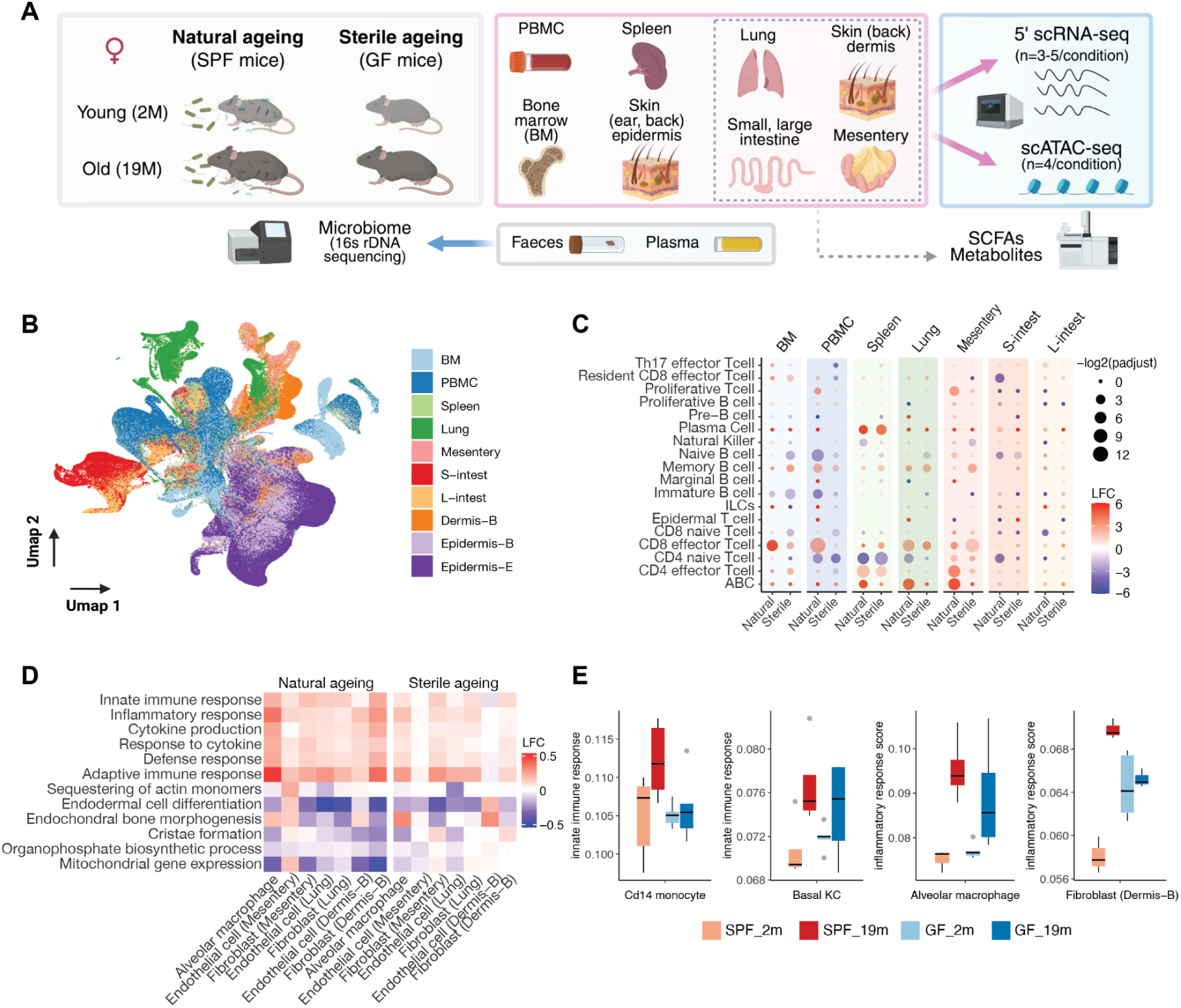
Single-cell analysis of ageing mouse tissues under SPF (natural) and GF (sterile) conditions (A) Schematic overview of the Mukin Mouse Ageing Atlas experimental design. (B) Integrated UMAP visualisation of scRNA-seq (180,296 cells) and scATAC-seq (289,489 cells) with single cells labelled by tissue origin. BM: bone marrow, S-intest: small intestine, L-intest: large intestine, Dermis-B: dermis from back skin, Epidermis-E: epidermis from ear skin. (C) Lymphoid cell type population changes in natural and sterile ageing across tissues. Cell proportions for each mouse replicate were first determined using scRNA-seq data for each tissue (n=3-5 per condition) to determine LogFCs between old (19m) and young (2m) mice for each tissue (see Figure S1Q for other cell types). Dot sizes are scaled by significance (-log2(p-adjust), see methods). (D) Gene ontology changes across a set of cell types at single-cell level resolution for natural and sterile ageing shows upregulation of inflammatory pathways in sterile ageing. Box plots of selected cell types showing distribution of gene ontology term scoring at pseudobulk averages (n=3-5).

Collectively, these findings highlight the ability of our approach to distinguish between intrinsic and microbiota-attributable ageing features.

We next asked whether age-associated changes are primarily dictated by intrinsic cell identity or by the tissue environment. We calculated the Sørenson similarity coefficient based on gene expression profiles for natural and sterile ageing separately across all cell types, followed by hierarchical clustering. In sterile ageing, clustering was predominantly driven by similar cell types, such as subpopulations of B cells and T cells, regardless of tissue origin (Figure S1R). In contrast, natural ageing showed clustering more by tissues rather than by cell types, grouping seemingly unrelated cell types, such as alveolar macrophages, endothelial cells, and fibroblasts in the lung (Figure S1R). Gene ontology analysis suggested the higher inflammatory state in natural ageing may drive such differences (Figure 1D-E). Notably, inflammation pathways were also elevated in sterile ageing, albeit to a lesser extent, a pattern indicative of sterile inflammation, that is, inflammation in the absence of infection or presence of microbiota. These observations suggest that cell identity is the primary determinant of age-associated transcriptome changes, while the tissue microenvironment exerts an increasingly prominent influence with age, consistent with recent observations of immune cells profiling in human tissues [24]. To further test this notion, we compared gene expression changes of keratinocytes (KCs) from two epidermal layers (basal and spinosum) across two skin regions (back and ear). Commonly upregulated genes were most enriched within the same tissue region (either back or ear) but markedly reduced across tissues or regions (Figure S1T and S1U). Taken together, these findings suggest that sterile inflammation is a hallmark of intrinsic ageing, and while cell identity fundamentally shapes ageing traits, tissue environment significantly modulates cellular states, particularly through inflammatory responses.

### Microbiota modulates cellular states during ageing

We sought to distinguish between cell types in which microbiota influences cellular states during ageing and those that age predominantly through intrinsic mechanisms. We reasoned that if microbiota substantially impacts cellular states during ageing, the correlation in gene expression changes between natural and sterile ageing would be low; conversely, a high correlation would suggest a predominantly intrinsic ageing process. To assess this, we calculated the Pearson correlation of age-associated gene expression changes between natural and sterile ageing for each cell type. Intestinal cell types, including intestinal stem cells, tuft cells, and enterocytes cells, exhibited the weakest correlations (Figure 2A), thereby indicating that their cellular states are strongly influenced by the presence or absence of microbiota during ageing. This finding aligns with the expectation that gut-resident cells would be particularly sensitive to microbial signals. Notably, CD16^+^ monocytes and pre-B cells also showed low correlations, thus suggesting microbiota-derived factors may regulate these cells. In contrast, alveolar macrophages, CD8^+^ naive T cells, hair follicle cells, and ABCs displayed high correlations between natural and sterile ageing, relationships that point to more intrinsically driven ageing processes in these cell types.

**Figure 2.**
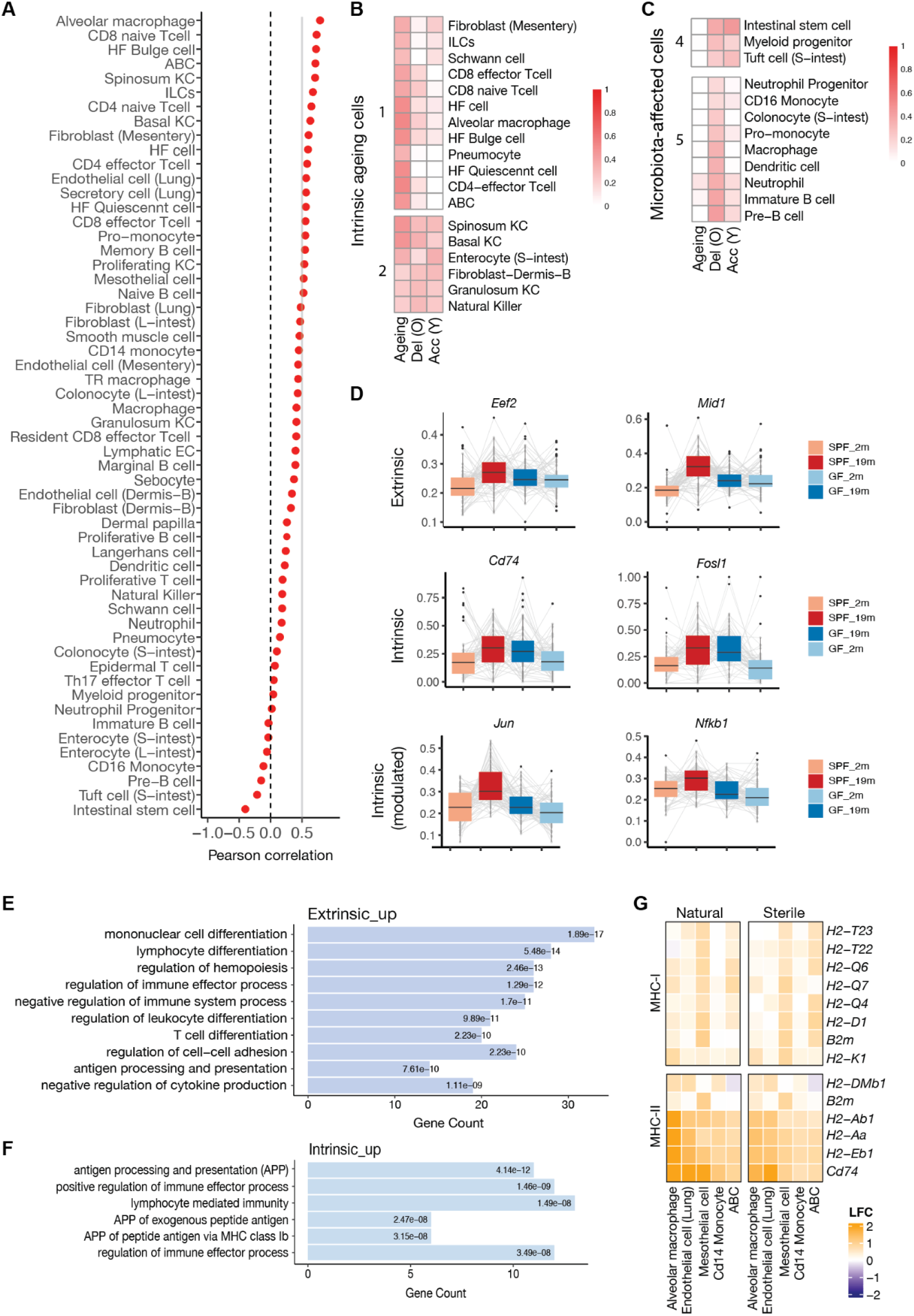
T**r**anscriptomic **profiling of microbiota effects during ageing** (A) Pearson correlation analysis between natural and sterile ageing for the listed cell types, showing the cell types that age more in an intrinsic (above 0.5 correlation) or extrinsic manner (negative correlation). (B-C) Effects of the GF condition on individual cell types. “Ageing” column shows similarity between natural and sterile ageing; (B) shows intrinsically ageing cell types with high similarity values and (C) shows microbiota-affected cell types with low similarity values. See Figure S2A for the full list of cell types. High similarity score in ‘Del (O)’ column suggests ageing is delayed by the GF condition, derived from similarity score between natural ageing DEGs and ‘old_GF vs old_SPF’ DEGs. ‘Acc (Y)’ column shows the extent of accelerated ageing by the GF condition based on similarity scores between natural ageing DEGs and ‘young_GF vs young_SPF’ DEGs. (D) Examples of upregulated intrinsic and extrinsic ageing genes. Average expression of indicated gene per cell type is visualised as a box plot for all cell types. Grey lines link the expression of the same cell type per condition. (E-F) Selected GO terms that are enriched in upregulated ageing-extrinsic (E) and intrinsic (F) genes. (G) Upregulation of MHC-I and MHC-II genes are one of the hallmarks of intrinsic ageing.

Next, we explored whether microbiota accelerates or delays ageing paths in the cell types they influence. Given the microbiota’s established roles in tissue homeostasis and impact on young animals, we also investigated whether the GF condition induces transcriptomic changes that resemble natural ageing in young mice. We assessed the similarity of gene expression changes by comparing differentially expressed genes (DEGs) between mice under SPF and GF conditions, using natural ageing DEGs as a reference. For each cell type, we performed pairwise comparisons to determine the effect of the GF condition: 1) during ageing (natural vs. sterile ageing), 2) in young mice (young GF vs. young SPF), and 3) in old mice (old GF vs. old SPF). DEGs were categorised based on the directionality of change relative to natural ageing. Genes changing in the same direction (for example, both upregulated) were assigned as “similar”, while those with opposing directions were categorised as “opposite”. We then calculated a Sørensen similarity score for these six gene sets relative to natural ageing DEGs, and grouped cell types by clustering (Figure S2A). The first column of the heatmap (“Ageing”) reflects *how similarly cells age* between natural and

GF ageing. Clusters 1, 2, 6, and 7 showed high similarity, suggesting these cell types age predominantly in an intrinsic manner (Figure 2B and S2A-B). Conversely, cell types in clusters 3, 4, and 5 showed low similarity, which shows the microbiota modulates their cellular states more during ageing. Intriguingly, while the GF condition had a delayed ageing effect on most cell types in old mice (positive similarity for “Del (O)” in Figure 2C), the GF condition appeared to accelerate ageing in many cell types in young mice (high similarity in “Acc (Y)” reflecting DEGs more closely resembling natural ageing genes, in Figure 2C and S2A). Such observation likely reflects the beneficial role of the microbiota in young, which is compromised with old age by dysbiosis. Overall, the GF condition impacted nearly all cell types to varying degrees, thereby highlighting the microbiota’s broad and context-dependent roles in shaping tissue homeostasis and cellular states during ageing.

Based on the idea that the most frequently dysregulated ageing genes across cell types and tissues would create microenvironments that make tissues susceptible to developing age-related diseases, we next seeked for potential genes and pathways that are frequently dysregulated across multiple cell types during ageing. DEGs were categorised into four groups for each cell type: intrinsic (dysregulated in the same manner between natural and sterile ageing), extrinsic (dysregulated in natural ageing only), sterile (dysregulated in sterile ageing only), and reversed (genes changing in opposite directions such as upregulated in one and downregulated in the other). We filtered for genes that are dysregulated in the same direction in more than three cell types (> +/-0.25 LFC) (Figure S2C, Supplementary table 6). The extrinsic category contained the most dysregulated genes, followed by sterile and intrinsic, results that reflect the substantial impact the microenvironment has on ageing cellular states, whereas reversed genes were rare (Figure S2D). Out of 88 genes that were found to be upregulated in an extrinsic manner, *Mid1* (encodes an E3 ubiquitin ligase, also known as *Trim18*) and *Eef2* (encodes a translation elongation factor) were the most dysregulated genes in this category, in 17 and 16 cell types, respectively(Supplementary table 6, Figure 2D). Amongst the intrinsic ageing genes, top three genes were of major histocompatibility complex class I (MHC-I) genes (*H2-K1, H2-D1* and *H2Q7*), upregulated in eleven cell types. *Cd74*, which encodes an invariant chain and is an essential component of the MHC class II (MHC-II) antigen presentation pathway, was also intrinsically upregulated in 8 cell types including non-immune cells such as endothelial, mesenchymal, and mesothelial cells. These observations suggest antigen presentation pathways in general are upregulated across multiple cell types. An AP-1 TF *Fosl1* gene was also upregulated in 8 different cell types in an intrinsic ageing manner (Figure 2D). Furthermore, we noticed several caspase genes, including *Casp1* were frequently dysregulated (Figure 2D, Supplementary table 6), which are critical components of the inflammasome. Inflammasomes are sensors of infections and sterile stressors such as cellular damage, and upon their activation produce activated Caspase-1. These activated Caspase-1 are essential in cleaving inactive precursors of IL-1β and IL-18 into their active forms that are potent innate cytokines and are critical to induce innate immune response [42], [43], [44]. We categorised a set of genes that were intrinsically dysregulated at significantly larger magnitudes in natural ageing than sterile ageing (“intrinsic-modulated”), which included another AP-1 TF family gene *Jun* and the NF-κB pathway gene *Nfkb1* that were upregulated (Figure 2D). Collectively, innate immune pro-inflammatory genes were frequently upregulated across cell types and tissues in an ageing-intrinsic manner.

We found some genes were upregulated in one cell type, and downregulated in another cell type. To identify robust genes that are dysregulated in the same direction across cell types, we took 269 extrinsic ageing genes that showed upregulation only across cell types (Figure S2E) for pathway analysis (Figure 2E). Immune-related functions were predominantly enriched, such as immune cell differentiation (Figure 2E, Supplementary table 7), while downregulated genes were enriched in protein synthesis-related processes (Figure S2F, Supplementary table 7). Pathway analysis of 55 uniquely upregulated intrinsic ageing genes in all dysregulated cell types showed enrichment in antigen presentation pathways (Figure 2F, Supplementary table 7). Indeed, upregulation of MHC-II genes and MHC-I genes to a lesser extent were particularly prominent (Figure 2G), which contrasted with their reduced expression in professional APCs, such as dendritic cells (Figure S2G, Supplementary table 7). In comparison, RNA splicing, protein folding, and translation pathways, all of which are known to be intrinsic cellular ageing features, were prominent in sterile-specific upregulated genes (Figure S2F, Supplementary table 7). Taken together, we have identified groups of cell types whose cellular ageing traits are affected or not affected by microbiota. Our analyses also highlight upregulation of pro-inflammatory genes and pathways as intrinsic ageing signatures across diverse cell types, dynamic ranges of which can be modulated by microbiota. As these signatures were as prominent in sterile ageing, we suspect these features likely arise from the accumulation of cellular stress and damage.

### Effects of microbiota on motif activity regulation during ageing

Transcription factors (TF) orchestrate gene expression programs that define cell identity and function, whose activity is regulated at both epigenetic and gene expression levels. To understand how ageing, both natural and sterile, alters these regulatory programs, we next profiled TF activity changes during ageing. Epigenomic TF activity was inferred from chromatin accessibility of TF binding motifs (hereafter referred to as motif activity), encompassing both promoters and distal regulatory regions such as enhancers [45]. In parallel, transcriptomic TF activity was estimated based on the expression of known target gene sets, or regulons (hereafter referred to as regulon activity) [46]. These two activity scores were derived at single-cell resolution to determine dysregulated (upregulated or downregulated) TFs within each cell type, and the number of cell types affected by each TF was quantified to identify frequently dysregulated TFs with ageing across cell types (Supplementary Table 8). Among the upregulated clusters (Figure 3A), cluster 10 included the most frequently dysregulated TF activities in an ageing-intrinsic manner. Notably, this cluster included well-established age-associated TFs including members of the AP-1 family (*Jun*, *Fos*, *Fosb*) and the NF-κB family (*Rela*, *Nfkb1*), results that underscore the robustness of our integrative analytical approach. We noticed that an increase in AP-1 family regulons, such as Jun and Fosb were higher in natural ageing compared to sterile ageing (Figure 3B). Together with our earlier observation of AP-1 genes themselves being identified as the ‘modulated intrinsic’ ageing genes (Figure 2D), these results suggest the upregulation of AP-1 TF-regulated genes is at least partially attributable to the microbiota. This is a novel finding as microbiota modulating the age-associated AP-1 TF upregulation has not been reported before. Interestingly, we identified highly dysregulated TFs whose motifs opened or closed in sterile ageing-specific manner (Figure 3A, cluster 5 for upregulation and Figure S3A-B, top cluster for downregulation). Sp/Klf (Specificity protein/Kruppel-like factors) family TF genes were particularly enriched in closed TF motifs in sterile ageing-specific manner, including one of the Yamanaka factors *Klf4*, raising an intriguing link between this family and the microbiota. Next, chromatin accessibility to gene expression correlation analysis for genes identified a set of intrinsically upregulated ageing genes, including *Cd74*, *H2-Q6*, *Casp1* and *Casp4* (Figure 3C, cluster 1, S3C-E, Supplementary Table 9). We found that the Caspase genes *Casp1*, *Casp4* and *Casp12* are located within the same topological domain and all became more accessible during ageing (Figure S3F), a pattern that suggests this region experiences age-associated chromatin conformation change. Additionally, a widely recognised marker of cellular senescence *Cdkn2a* gene, which encodes the cyclin-dependent kinase inhibitor 2A (p16^INK4a^), was frequently dysregulated across multiple cell types at the chromatin accessibility level only (cluster 2 in Figure 3C-D). Such alternation in epigenetic state without detectable upregulation of gene expression may reflect the non-constitutive nature of p16 expression, and likely highlights the limitation of relying only on transcriptomic data to identify senescent cells. Collectively, we found upregulation of AP-1 family-regulated genes were attenuated in the germ-free condition via epigenetic changes and identified TF motifs that are dysregulated in sterile ageing specific-manner. Overall, many TFs showed dysregulated motif activities without corresponding regulon dysregulation. It is unknown whether such regions represent compromised chromatin structure maintenance or epigenetic ‘memory’ states, in a similar manner to those observed in trained immunity and inflammation memory [47], [48].

**Figure 3.**
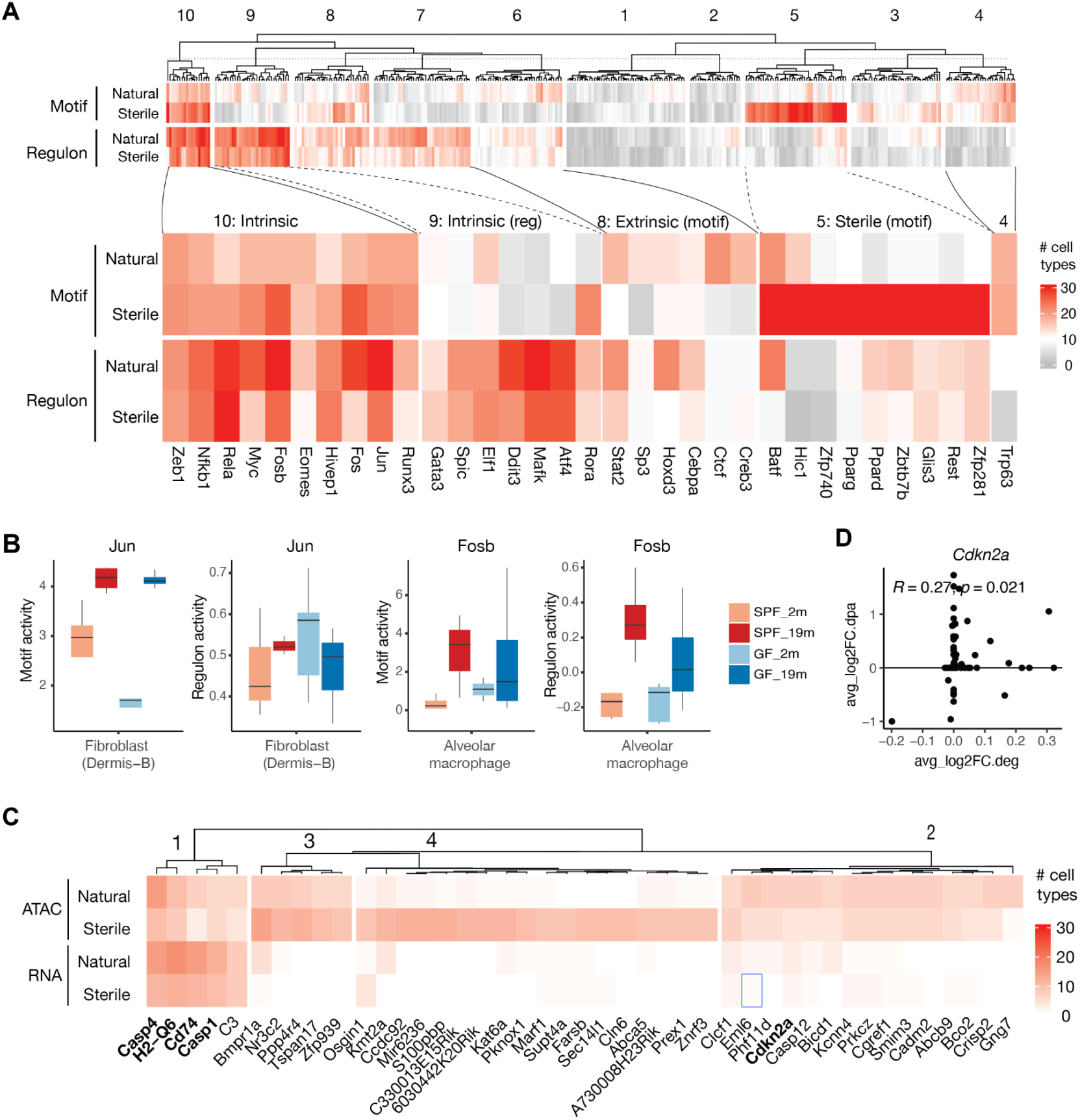
E**f**fect **of microbiota on motif activities during ageing** (A) Number of cell types that showed upregulated TF motif and regulon activities during natural and sterile ageing. TFs are grouped and clustered based on hierarchical K-means clustering and dendrogram clustering, respectively. Heatmap shows number of dysregulated cell types. Selected TFs from clusters are highlighted. (B) Motif and regulon activities for AP-1 TFs *Jun* and *Fosb* for indicated cell types showing higher increase in natural ageing compared to sterile ageing. Box plots display scores of pseudobulk averages (n=3-5). (C) Groups of genes that showed frequent chromatin opening with age (Figure S3E for chromatin closing genes) across multiple cell types. Heatmap shows the number of dysregulated cell types. Genes are grouped and clustered based on hierarchical clustering and dendrogram clustering, respectively. Gene accessibility LFC (y-axis) and expression LFC (x-axis) for *Cdkn2a* gene are plotted for all cell types. Calculated Spearman correlation is shown.

### Alveolar macrophages senescence is accelerated by dysbiosis

We noticed alveolar macrophages consistently showed a high magnitude of age-associated changes compared to other cell types across analyses thus far. To identify the source of such prominent changes, we carried out differential analysis between old and young alveolar macrophages, which revealed substantial increases in both chromatin accessibility and gene expression of cytokines, NF-κB-related genes, and MHC-II genes in both natural and sterile ageing (Figure 4A, S4A). As many of these genes are implicated in senescence, we next assessed the degree of senescence in alveolar macrophages relative to other cell types with an enrichment analysis with a compiled senescence-associated gene set termed SenMayo [49]. Alveolar macrophages had the highest senescence score during natural ageing, followed by memory B cells and endothelial cells (Figure 4B, S4B). Immunostaining of lung tissues with senescence marker SA-β-GAL with alveolar macrophage marker Siglec-F, along with other senescence characteristics such as expanded cellular volume and reduced sphericity, confirmed more frequent senescent alveolar macrophages in old SPF lungs compared to the young (Figure 4C). Elevated senescence scores were also observed with publicly available ageing lung datasets [23], [36] for alveolar macrophages, albeit at a lower level (Figure S4C). Further, the senescence score for alveolar macrophages was considerably higher in natural than sterile ageing, both at chromatin accessibility and gene expression levels, a contrast which was attributed to more mice harbouring senescent alveolar macrophages (four SPF mice compared to two GF mice, Figure 4D, S4D). Thus, alveolar macrophage senescence appears to be an ageing-intrinsic feature that the microbiota modulates. Such difference in senescence frequency, however, was not observed in other cell types, such as skin dermal fibroblasts in the same mice (Figure S4E-F), suggesting the extent of senescence varies by cell type and tissue context.

**Figure 4.**
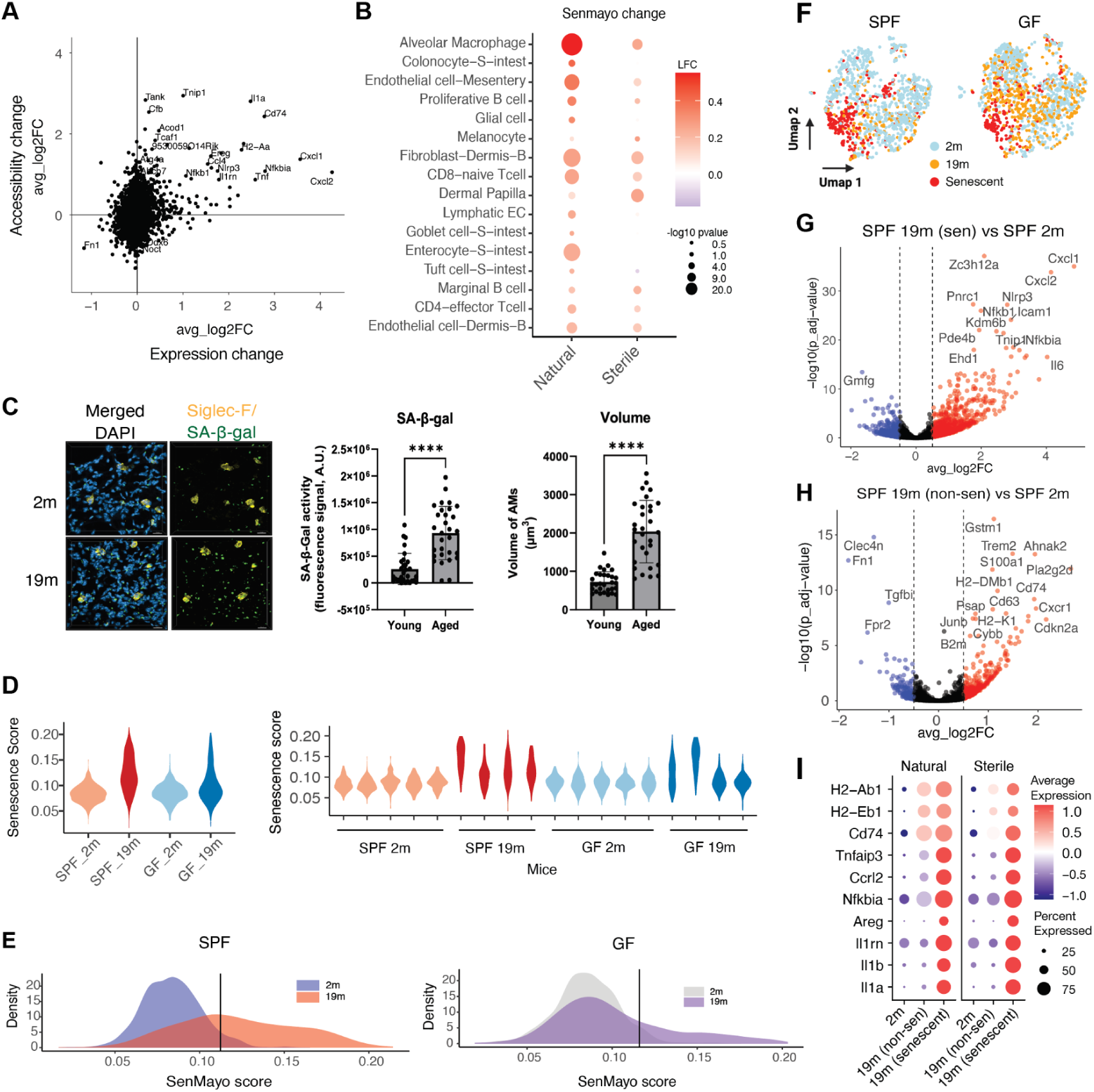
S**t**erile **conditions attenuate alveolar macrophage senescence** (A) Correlation of age-associated chromatin accessibility to gene expression changes in alveolar macrophages from SPF mice (GF mice in Figure S4A) shows many pro-inflammatory genes are highly upregulated. For each peak, a gene was chosen based on the most accessible promoter peak. Scores are shown as LFC. (B) Senescence score increases with age across cell types, most prominently in alveolar macrophages in natural ageing (see Figure S4B for the full list of cell types and sterile ageing). Mean senescence score was calculated with AUCell with SenMayo geneset. Significance was tested with Wilcoxon rank-sum test (P value <0.05). (C) Immunofluorescence image of young (2 months) and old (19 months) lung tissues, stained for DAPI (blue), Siglec-F (yellow; AM marker) and SA-β-GAL (green; senescence marker) (left). Quantification of signals (middle) and cellular volume (right) shows an increase in senescent alveolar macrophages in old lungs. (D) Senescence score distribution of all alveolar macrophages per condition (left) and per mouse (right). (E) Senescence score distribution of alveolar macrophages in SPF (left) and GF (right) mice. Vertical separation line denotes the 95th quantile based on senescence score of young alveolar macrophages, above which are classified as senescent-like. (F) UMAP visualisation of alveolar macrophages labelled by age and state. (G-H) Volcano plots show DEGs of alveolar macrophages between senescent (sen) and young (G) and non-senescent to young (H). Red: upregulated, blue: downregulated. A dotplot of selected genes showing upregulation of both ageing (top three) and senescent-associated genes are upregulated in senescent alveolar macrophages. Dot size indicates percentage of cells expressing indicated gene. Mean expression per group is shown by a red-to-blue gradient.

A wide distribution of senescence score as well as SA-β-GAL staining suggested alveolar macrophages from old mice comprised a mixture of senescent and non-senescent subpopulations. To better characterise the senescent population, we defined senescent alveolar macrophages by applying a 95th percentile cutoff score to young alveolar macrophages (Figure 4E-F). We next performed pseudobulk DEG analysis between senescent and non-senescent subpopulations. IL-1 family cytokines (*Cxcl1*, *Cxcl2*, and *IL1b*) and NF-κB pathway (*Nfkbiz*, *Tnf*, *Nfkbia*, and *Tnfaip3*) genes were upregulated in senescent alveolar macrophages (Figure 4G-H, S4G, Supplementary Table 10), confirming their inflammatory state. We noticed downregulation of an extracellular matrix *Fn1* gene between non-senescent alveolar macrophages from old mice to young (Figure 4H, S4I, Supplementary Table 10) that encodes for a glycoprotein called fibronectin-1 (FN1). As it is thought to play a role in tissue repair [50], *Fn1* downregulation may be an important contributing factor in age-associated tissue repair decline in the lung. We also observed elevated expression of previously identified “intrinsic ageing genes”, such as *Cd74* and MHC-II genes *H2-DMb1*, *H2-Eb1* and *H2-Ab1*, in both senescent and non-senescent alveolar macrophages from old lungs (Figure 4H-I, S4H-I,Supplementary Table 10). Little differences were detected between senescent alveolar macrophages from SPF and GF mice (Figure S4J).

To explore the functional state of senescent alveolar macrophages, we next used scRNA-seq data to predict their metabolic state, specifically via the expression of 518 metabolic enzymes across 22 central metabolic pathways [51]. Senescent alveolar macrophages had large age-associated reductions in predicted lactate uptake and ATP production functions (Figure S4K). Consistently, total lactic acid levels rose in old mouse lungs during both natural and sterile ageing (Figure S4L), although how much this change stems from alveolar macrophage senescence state remains to be determined. Lactate was long considered to be a mere byproduct of glycolysis, however, it is now evident that lactate level influences cellular functions by acting as a metabolic switch as well as epigenetic gene regulation through recently discovered histone lactylation [52], [53], [54]. In light of our observation, senescent alveolar macrophages may contribute to age-associated increase of lactic acid in the lung, however its functional consequence on the tissue remains unknown. In summary, presence of microbiota accelerated age-associated alveolar macrophage senescence, which is accompanied by a metabolic state change that may contribute to an elevated total lactate level within the lung. Further investigations are needed to better understand what factors in the lung drive senescence of alveolar macrophages possibly at a faster pace compared to other cell types.

### Emergence of ABCs is a prominent feature of intrinsic ageing

ABC expansion was the most striking intrinsic ageing feature we identified in this atlas, showing no difference in magnitude of increase between natural and sterile ageing conditions across tissues (Figure 1C). ABCs are known to accumulate in healthy ageing, infections and vaccination, and play a role in effective clearance of viruses upon infections [28], [29], [55], [56], [57]. However, they are also linked to pathogenicity in autoimmune diseases such as systemic lupus erythematosus (SLE), multiple sclerosis, and rheumatoid arthritis [29], [58], [59]. In-line with our finding, ABC expansion has been reported in the blood of old GF mice, more prominently in female mice [60]. Amongst all B cells from the atlas, we identified eight transcriptionally distinct clusters: pre-B (expressing *Vpreb1*), immature (*Fcer2a* negative), proliferative (expressing *Mki67*), B2 (expressing *Fcer2a*), marginal (expressing *Cr2* high), memory (expressing *Zbtb32*), ABC expressing *Cd11c* and *Tbx21* (encodes for Itgax and T-bet, respectively), and plasma cells (expressing *Sdc1*) (Fig 5a, S17A) [61]. We identified *Fcer1g* expression as the most robust marker distinguishing ABCs from other B cell populations, with stronger and more consistent expression than previously reported markers such as *Zbtb32* and *Tbx21* (Figure 5B-C). *Fcer1g* encodes the Fc receptor γ-chain (FcRγ), a component of innate immune signalling, supporting the concept that ABCs exhibit innate-like characteristics. Increased chromatin accessibility at *Tbx21* and *Fcer1g* genes were observed in ABCs (Figure S5B-C). ABC expansions were detected in multiple tissues, including the bone marrow, PBMC, spleen, lung, and mesentery (Figure 5D). We detected little tissue-specific gene expression profiles (Figure S5D) suggesting while ABCs may adapt subtly to local environments, they largely reflect a circulating, tissue-independent cell state. We did not detect large differences between natural and sterile conditions (Figure S5E). Consistent with our earlier observation, regulon activities of AP-1 factors *Jun* and *Fos* were higher in ABCs from natural ageing compared to sterile ageing (Figure S5F, Supplementary Table 11). A small elevation of inflammatory factors *Il4* and *Il6* at both chromatin accessibility and gene expression were observed in natural ageing ABCs (Figure 5E, S5H). There have been little prior reports regarding IL-4 production by ABCs, and whether its functional role may be more in autoimmunity or B cell differentiation into plasma cells, remains to be determined.

**Figure 5.**
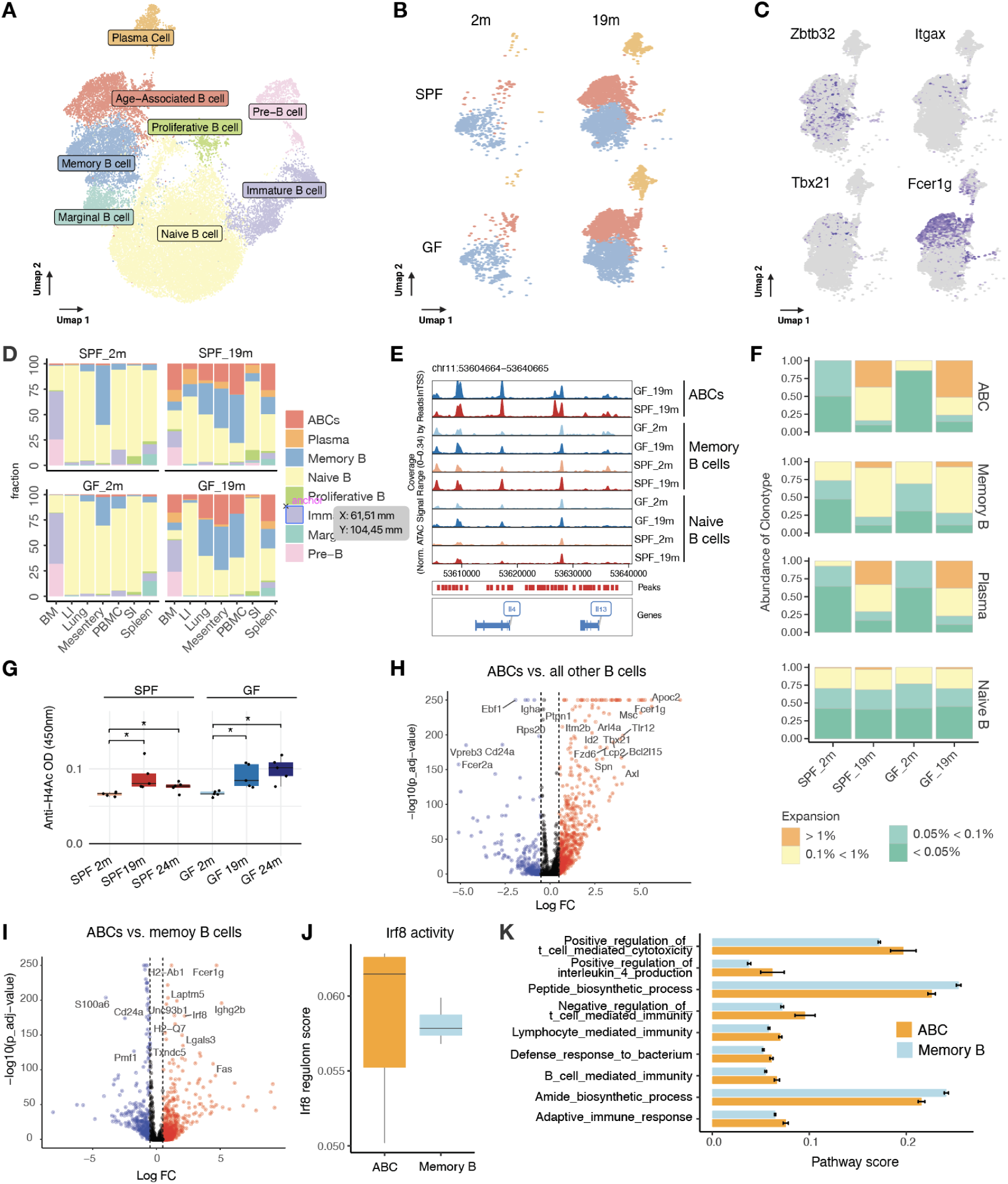
A**B**Cs **intrinsically expand with age** (A) UMAP visualisation of all B cells in the Mukin atlas. (B-C) UMAP visualisation of plasma cells, memory B cells, and ABCs per condition (B) and ABC markers (C). (D) Frequency of B cell subpopulations across tissues and conditions shows ABCs increase with old age across tissues. (E) Genome browser showing chromatin accessibility of *Il4* locus is more open in ABCs. (F) Clonality analysis shows an increase in similar clones for ABCs and plasma cells with old age. Occurrence is calculated per clonal type; degree of expansion is depicted by binning into >1% (orange), 0.1% < 1% (light green), 0.05% < 0.01% (green), and < 0.05% (dark green). (G) An ELISA assay shows an increase for anti-H4Ac autoantibody in plasma with old age. n=5; t-test; *P value < 0.05. (H-I) Volcano plots of differentially expressed genes in ABCs compared to all other B cells (H) and to memory B cells (I). Red: Upregulated, blue: downregulated. (J) Irf8 regulon score is higher in ABCs compared to memory B cells. Selected enriched gene ontology terms of significantly upregulated genes (p adjust<0.05) between ABCs (orange) and memory B cells (blue) are shown.

We next carried out clonal type analysis of B cell receptor (BCR) repertoire by Trust4 (version 1.0.7) [62], which strongly suggested ABC expansion is driven by clonal expansion (Figure 5F, S5I-L). Furthermore, consistent with the notion that ABCs are IgM-expressing memory B cells that differentiate into antibody-secreting plasma cells upon secondary antigen challenge [63], ABC and plasma cell populations exhibited clonal overlapped (Figure S5M).

Earlier reports showed that ABCs produce autoantibodies *in vitro*, including anti-chromatinIgG antibodies in response to TLR7 agonists and anti-dsDNA in a mouse lupus model [29], [64], thus we next checked autoantibodies levels in the plasma by ELISA. Indeed, we found anti-chromatin autoantibodies (against acetylated histone H4) increased during both natural and sterile ageing, whereas autoantibodies against dsDNA were not at detectable levels (Figure 5G). Other DEGs between ABCs to B cells identified *Apoc2* (associated with lipid metabolism) upregulation and *Ebf1* (a B cell TF) downregulation along with its regulatory gene *Vpreb3* (Figure 5H), changes that indicate a shift in B cell identity or state.

As ABCs are considered a type of memory B cells formed under inflammatory conditions [65], we next set to determine transcriptomic differences between ABCs and memory B cells, which showed *Fcer1g*, *Ighg2b*, and *Irf8* upregulation, along with strong downregulation of *S100a6* (Figure 5I). As IRF8 is required for B cell differentiation including class switching and maintenance of B cell identity [66], [67], [68], its upregulation as well as its activity (upregulation of its target genes; Figure 5J) likely reflects ABC activation as previously reported [57]. Curiously, pathway activity analysis of ABCs revealed enriched T cell-mediated toxicity as well as upregulated IL-4 production pathways (Figure 5K), which require further validation and remains to be seen what they entail. Taken together, ABCs intrinsically expand with age systemically through clonal expansion, with no signs of modulation by microbiota. How much they may contribute towards sterile inflammation remains to be determined.

## Discussion

By comparing natural and sterile ageing, we identified microbiota-dependent and -independent age-linked alterations. Effects of microbiota during ageing were context-dependent; overall their absence had a negative effect on many cells in young mice resembling premature ageing, whereas ageing-related expression changes were attenuated in old mice. This contrast likely comes from age-associated dysbiosis, whereby beneficial roles of the microbiota on tissue homeostasis become more pathogenic with age-induced dysbiosis. AP-1 TFs upregulation is one of the prominent features of ageing, also implicated in senescence [69], [70], which we found to correlate with microbiota presence. To our knowledge, a direct connection between ageing-associated dysbiosis and AP-1 has not been established, making this association a key novelty of our work. As a possible mechanistic insight, we found butyrate and propionate - microbiota-derived metabolites - increase with ageing, which are potent activators of AP-1 signaling *in vitro* [71], likely contributing to the attenuation of AP-1 activation in sterile ageing. An interesting prospect on the function of AP-1 lies within its epigenetic memory function, and recent studies have shown that *Fos* expression in astrocytes plays a critical role in memory recall [72]. We also determined ageing under SPF conditions resulted in more mice with alveolar macrophages in senescence, consistent with links between gut microbiota and senescence [73]. Inflammation is likely central to driving senescence in alveolar macrophages; however, the exact mechanism warrants further study. Notably, transcriptomic senescence signatures were higher in our dataset than in publicly available ageing lung resources, a variation that may reflect differences in environmental exposures (for example, air quality) or diet/chow composition. Despite efforts to identify senescent cells from scRNA-seq data [74], inferring senescence from transcriptomes remains challenging, as inflammatory and senescence gene sets overlap extensively, thus blurring distinctions between highly inflamed and truly senescent states. We therefore applied senescence score thresholding in an attempt to separate ‘inflamed but not senescent’ from senescent cells, and future work may determine an inflammation threshold that pushes cells into senescence.

Microbiota presence during ageing affected not only cells in direct contact with microbes, such as intestinal stem cells, tuft cells, and colonocytes, but also circulating myeloid cells ranging from progenitors to differentiated neutrophils, dendritic cells, and macrophages. These findings align with reports that microbiota-derived molecules regulate haematopoietic stem cell functions via the gut–bone marrow axis, such as microbiota-derived lactate in the activation of haematopoiesis and erythropoiesis in the bone marrow in a lactate receptor-dependent manner [75], [76]. Such differences observed between natural and sterile ageing in distant organs from the gut are likely due to microbiota-derived molecules such as short chain fatty acids and lipids that are transported by portal blood and lymphatic circulation, including by modes such as extracellular vesicles. It would be interesting to evaluate whether the microbiota-dependent metabolite sulfonolipid, which we have previously shown to elevate in multiple organs such as the spleen, kidney, and lung during ageing [77], contributes to epigenetic or other microbiota-dependent cellular ageing changes. We also identified *Mid1* gene that is upregulated in a microbiota-dependent manner with age. As inhibition or depletion of Mid1 results in downregulation of the mTORC1 signalling pathway, and mTOR inhibitor rapamycin results in lifespan extension [78], [79], further studies should elucidate whether intrinsic upregulation of *Mid1* is responsible for the mTORC1 signalling upregulation. Our study showcases the wide impact microbiota has on the epigenetic landscape not only in the gut but also in distant organs. Butyrate has long been known as a histone deacetylase (HDAC) inhibitor, and more recently discovered to act as a post-translational modification including histone lysine residues on H3K18 and H4K12, along with propionylation, and succination [80], [81]. Additionally, we observed a striking enrichment of the Sp/Klf family TF motifs closing in sterile ageing-specific manner. Some members such as Klf4 gene expression [82] and Sp1 acetylation have been linked directly to butyrate [83], raising a curious possibility that the entire family of Sp/Klf TFs could be regulated by SCFAs.

Microbiota-independent traits we identified included increased sterile inflammation and expanded ABCs, likely driving inflammageing. We found both MHC class I and II genes were upregulated intrinsically across cell types, a change reminiscent of interferon-gamma-driven inflammatory activation in the absence of microbial infections. Supporting this observation, we saw elevations in CD8^+^ effector T cells (which are restricted to MHC-I) and CD4^+^ effector T cells (which are restricted to MHC-II) in both natural and sterile ageing. Other studies have reported similar ageing-associated upregulation of MHC-II genes in various tissues and cell types [84], however we found it to be more of a systemic ageing phenomenon than to specific cell types. Consistent with our observation that it is intrinsic to cellular ageing, a study used intestinal organoids shielded from extrinsic signals from the microenvironment and showed cell-intrinsic inflammageing (MHC-II upregulation) persists in intestinal stem cells in culture [84]. Moreover, this cell-intrinsic inflammageing is primed at the chromatin level, and thus authors suggest memory of inflammation state is retained in the chromatin. Intriguingly, one of the MHC-II-related genes, *Cd74*, was among the most strongly upregulated intrinsic genes in our dataset and is also frequently upregulated across various cancer types [85]. Tumours have been reported to express MHC-II genes, which are associated with higher prognosis [86]. While visible solid tumours were excluded during sample preparation, it remains to be seen to what extent these findings reflect early tumourgenic changes versus a pre-disease state cells reach during ageing that makes them susceptible to developing cancer and other age-related diseases.

The underlying mechanism of sterile inflammatory activation we have identified as intrinsic to ageing likely involves the Nlrp3 inflammasome. For more than a decade, sterile inflammation has been proposed as a major driver of inflammageing, primarily via host-derived damage-associated molecular patterns (DAMPs) [87], [88]. Ageing-associated DAMPs induce chronic activation in tissue-resident macrophages such as alveolar macrophages, microglia in the brain, and Kupffer cells in the liver, leading to inflammatory damage and tissue dysfunction [89]. Here, we showed chromatin accessibility of *Caspase-1* and *Caspase-4* become open and are upregulated with old age in a striking number of cell types including immune and non-immune cells. Together with our finding that ABC expansion occurs during sterile ageing to the same extent to natural ageing raises the possible contribution of ABCs towards sterile inflammation, and therefore inflammageing. This idea is in-line with our finding that ABCs express the innate marker *Fcer1g* and marks innate-biased programs. It is of note that ABC expansion was particularly prominent in our atlas, likely because female mice were used. Autoimmune diseases are more prominent in females, owing to a higher *Tlr7* gene expression as a result of escaping X inactivation in immune cells, including ABCs [90]. TLR7 stimulation is one of the key components of ABC activation, likely explaining a more prominent ABC expansion in female mice [29], [57], [60], [91]. Further investigations are needed to determine to what extent ABCs contribute towards inflammageing, as well as identifying sources of ABC activation *in vivo*, including DAMPs, food antigen and transposable element transcripts such as LINE1 elements, known to act as sources of inflammation [92].

Although we did not assess intestinal permeability in this study, it has been shown to increase with ageing-associated dysbiosis and likely contributes to the microbiota-dependent changes we have identified. One of the limitations of our study is that GF mice are not educated by microbiota, making it difficult to disentangle early life colonisation developmental effects that involve immune benefits from ageing-related changes in adulthood. Future experiments, such as administering antibiotics from young adulthood and exposing old GF mice to infections or other stress conditions, should provide further insights. Furthermore, to keep mice under GF condition, cages were placed in isolators that also filtered out air pollutants and impurities, thus we cannot rule out the possibility that acceleration of alveolar macrophage senescence could come from differences in the air quality without further investigations. Given its substantial rise during ageing, minimising sterile inflammation is likely a key to enabling healthy longevity, as a higher inflammatory state broadly impairs cellular function. While animal models are maintained in controlled, relatively clean environments that may not mirror human lifestyles, our extreme-condition functional atlas nonetheless delineates ageing features modulated by an extrinsic factor, namely microbiota, versus intrinsic aspects that accrue with time. Our study illuminates, for the first time, how cells age in tissues under germ-free conditions. Our single-cell resolution at both the epigenomic and transcriptomic levels demonstrates, in the most comprehensive manner to date, that sterile inflammation and ABC expansion are major intrinsic ageing traits, likely to be prominent drivers of inflammageing. We expect the Mukin Mouse Ageing Atlas to serve as a resource for uncovering mechanisms of cellular ageing in currently underexplored contexts, such as microbiota-dependent epigenetic regulation. More broadly, identifying microbiota-dependent circulating metabolites, such as lipids and short chain fatty acids that circulate to distant organs to tune epigenetic and therefore cellular states, may serve as targets for interventions that preserve resilient epigenomes that are less susceptible to developing diseases.

## Methods

### Mice

All experimental procedures were approved and performed in accordance with the Institutional Animal Care and Use Committee of the RIKEN Yokohama Campus (2019-015(2)). Four-week-old female C57BL/6N mice were purchased from CLEA Japan Inc. (Shizuoka, Japan). GF mice were housed and maintained in GF conditions in vinyl isolators with an alternating 12-h light/dark cycle at 23 °C with free access to food and water at the animal facility at the RIKEN Yokohama campus. All mice were fed AIN-93M chow (CLEA Japan, Tokyo, Japan). The GF condition was validated by using qPCR for bacterial count in faeces. Collected tissues that had tumours or abnormal phenotypes were avoided as much as possible for single-cell analysis.

### Tissue collection & single-cell dissociation

Single-cell dissociation was performed upon tissue harvesting as detailed below. Tissues were then cryo-preserved in CELLBANKER 1 (Zenyaku #11910) for subsequent scRNA-seq and scATAC-seq library preparation (Figure 1A and Table S1). Tissues from the same mice (three to five mice depending on tissues) were processed for single-cell data production.

#### Bone marrow

Hindlimbs were removed above the hip joint. The skin was teased away and separated in one piece from the leg. The foot was detached from the leg below the calcaneum. Scissors were used to clear flesh away from the bones. Next, bones were washed with 2 mL of Hank’s Balanced Salt Solution (HBSS) (Sigma #H9394) and the rest of the flesh was removed with a Kimwipe (Fisher Sci #06-666). Upper and lower legs were separated from each other at the patella. The fibula was disconnected from the tibia and discarded. A 5 mL syringe was filled with wash solution (HBSS (Sigma #H9394) + 2% FBS (Gibco #26140079) (ice cold)) and capped with a 23.5-gauge needle. The ends were cut off the bone to expose the bone marrow (red). The syringe was placed into the bone cavity and gently depressed to expel the wash solution. Tubes were spun in a refrigerated centrifuge at 4 degrees at 240g for 5 min. The pellet was resuspended in 1 mL RCL buffer (0.15M NH4Cl 0.01M Tris HCl Make up to 500 mL with RO Water; modified pH to 7.5 ±0.2) and incubated at room temp for 5 min. 5 mL FBS was added to neutralize. The sample was spun and washed with wash solution. Samples were resuspended in freezing media (90% FBS (Gibco #26140079), 10% DMSO (Nacalai #13408-64)) and filtered (Corning #352235). Samples were aliquoted into cryovials and stored at -80° C.

#### PBMC and Plasma

Blood was collected via cardiac puncture (using a 23-25 gauge needle) into a heparin-containing tube. The sample was spun at 1000g (RT) for 10 min. Plasma (supernatant) was collected, aliquoted, and frozen. 1 mL 30% Percoll was added to the remaining sample to a final volume of 10 mL. The sample was then vortexed and mixed well. A 2 mL 70% Percoll cushion was layered on the bottom of the tube, bringing the final volume to 12 mL. The sample was spun at 800g for 25 min (RT) with both the accelerator and brake off. The top layer was discarded and the middle layer collected into a fresh tube. The sample was washed with RPMI (GIBCO #11875085), then centrifuged at 800g for 5 min at 4 degrees. Cells were resuspended in CELLBANKER 1 (Zenyaku #11910) and stored at -80°C.

#### Spleen

The spleen was dissected whole from the mouse and placed in a petri dish with HBSS (Sigma #H9394). Using the bottom of a 5 mL syringe (Terumo #1SS05LE), the spleen was ground until it burst. The cell-containing medium was then filtered through a 40 μm cell strainer (Corning #352340). Cells were centrifuged at 300g for 5 min. The pellet was resuspended in 1 mL RCL buffer (0.15M NH4Cl 0.01M Tris HCl Make up to 500 mL with RO Water; modify pH to 7.5 ±0.2) and incubated at room temp for 5 min. 5 mL of FBS were added to neutralise (Gibco #26140079), and the tube was centrifuged at 300g for 5 min. The pellet was washed in HBSS, resuspended in CELLBANKER 1 plus (Takara Bio #CB021), and stored at -80° C.

#### Lung

The whole lung was dissected from the mouse and processed for single-cell dissociation as described previously in [93] with the modification at the end; the cell pellet was resuspended in 5 mL 30% Percoll (1.5 mL Percoll (Merk #P164), 3.5 mL HBSS (Sigma #H9394)) and centrifuged at 2000 rpm for 25 min at room temperature. The pellet was then washed in lung medium (7 mL RPMI, 50 μm /nL Liberase™, and 10 μg/mL Dnase), centrifuged at 1500 rpm for 5 min at 4° C, and suspended in 3 mL of the lung medium for cell counting.

#### Mesentery

Mesentery was removed and cut into two pieces, one of which was used for single-cell dissociation. The adipose tissue was first chopped and minced by scissors on the inside wall of a 15 mL Falcon tube containing 7 mL of cold 4% BSA DMEM, making sure the tissue did not dry out. Minced pieces were then placed into the solution and mixed by inverting the tube upside down several times, ensuring all fat pieces were submerged in the solution. 50 μL of Liberase DH and 0.7 μL of DNaseI were added to each tube and incubated at 37° C until warm. The solution was then transferred to a shaker set at 37° C, 150 rpm for 60 min, with one manual shake at 30 min and another manual shake after 60 min. Volume was then topped up to 15 mL with 10% FCS RPMI and centrifuged at 1500 rpm for 5 min at room temperature. The supernatant and fat layers were removed with an aspirator, the pellet was dislodged by tapping, and 10% FCS RPMI was added to make up to 15 mL. The solution was then passed through a piece of 37 μm mesh and centrifuged. After removing the supernatant, 1 mL of ACK Lysing Buffer was added and incubated for 2 min for red blood cells lysis. 10% FCS RPMI was added to bring the volume up to 15 mL, mixed by tapping, passed through a piece of 37 μm mesh, and centrifuged. Supernatant was removed and 10% FCS RPMI was added to make 0.5 mL or 1 mL, depending on pellet size. Cells were counted and adjusted to be 1 x 1-^6 cells/mL with 10% FCS RPMI.

#### Small and large intestine

One-third of the length (∼15cm) of either the entire small intestine, including Peyer’s patches on the terminal ileum side, or the large intestine, excluding cecum, was dissected from each mouse. After eliminating intestinal-associated fat tissue, intestines were washed with the cold RPMI-1640 medium (Sigma Aldrich) to remove luminal contents and debris. Intestinal tissues were cut into smaller pieces and digested with 1.0 mg/ml Collagenase (Wako, Cat# 032-22364) for the small intestine or with 1.0 mg/ml Type 1A Collagenase (Sigma Aldrich, Cat# C9891) for the large intestine, then suspended in RPMI-1640 medium containing 2% foetal bovine serum (FBS) and incubated at 37° C for 15 min. The resultant supernatants from the collagenase digestion were collected and passed through a 100 μm cell strainer (BD Biosciences) after three cycles of these steps. The dissociated cells were collected after spinning down and passed through a 40 μm cell strainer (B.D. Biosciences). The single-cell suspension was diluted to 1x106 cells with cold 10% FBS in RPMI-1640 medium.

#### Skin (Back)

The dorsal epidermal and dermal cell suspensions were prepared from shaved back skin as previously described (Tsutsui et al., 2021). Briefly, the dermal adipose layer of dissected skin was physically scraped off with a scalpel. The skin tissue was treated with 0.25% trypsin (Nacalai Tesque) and 2 U/mL DNase I solution, epidermis-side up, at 37° C for 1 hour. The epidermal tissue was separated from the dermal tissue with a scalpel. To prepare the epidermal cell suspension, the separated epidermis was minced with scalpels and mixed well with repeated pipetting, followed by passing through a 40 μm cell strainer (BD Falcon). The remaining dermal tissue was minced and treated with 2 mg/mL collagenase type I (Gibco), 2 U/mL DNase I, 5 μg/mL hyaluronidase (Sigma) in HBSS with Ca and Mg (Gibco) at 37° C for 2 hours with gentle mixing. Single-cell suspension of dermal cells was obtained by repeated gentle pipetting and passed through a 40 μm cell strainer. Aliquots of single-cell suspensions were stored at -80°C in Bambanker (Nippon Genetics) until use for scRNA-seq.

#### Skin (Ear)

Epidermal keratinocytes from mouse ear were isolated as previously described with a slight modification (Usui et al., 2019). Dissected mouse ear skin were split into dorsal and ventral halves and put on 50 µL of 10 mg/mL dispase II (Roche, Indianapolis, IN) dissolved in 50 mM HEPES (pH 7.5) containing150 mM NaCl. After incubating at 37° C for 15 min, the epidermal sheets were carefully separated from the dermis. After rinsing with PBS containing 1mM EDTA, the epidermis was floated on 1 mL of Trypsin/EDTA solution (Kurabo Industries Ltd., Osaka, Japan) in a 35-mm plastic dish and incubated for 5 min at 37° C. Then, keratinocytes were separated mechanically by fine tweezers. Keratinocyte suspension was added with 1 mL of Trypsin Neutralizing solution (Kurabo Industries Ltd.) and passed through the cell strainer of a Falcon 5 mL Round Bottom Polystyrene Test Tube, with a Cell Strainer Snap Cap (Corning), then immediately diluted with 8 mL HuMedia-KG2 (without antibiotics) (Kurabo Industries Ltd.) and centrifuged. The cell pellet was resuspended in Bambanker and cryopreserved (Nippon Genetic Co. Ltd., Tokyo, Japan).

### Single-cell library preparation and sequencing

#### scRNA-seq (n=3-5 mice per condition)

Single-cell RNA-seq libraries were prepared according to the 10X Genomics user guide ‘Chromium Next GEM Single Cell V(D)J Reagent Kits v1.1’, except for the following modification from post-cDNA Amplification Reaction Cleanup. After 0.6x size selection, the supernatant was kept as small fragments, which we suspected may contain regulatory elements. 20 μL (0.8x) SPRIselect reagent was added to this supernatant and incubated for 5 min at room temperature. The beads were washed and eluted in 20 μL Buffer EB, which is referred to as ‘No Fragmentation short size cDNA’. Adapter ligation mix was prepared according to the manufacturer’s instructions with the following change; 10 μL of No Fragmentation short size cDNA was added to the ligation mix, to bring the total reaction volume to 50μL. The sample was returned to the thermocycler and the manufacturer’s protocol was resumed.

#### scATAC-seq (n=4 mice for all conditions)

Nuclei for the single-cell ATAC libraries were prepared according to the 10X Genomics demonstrated protocol ‘CG000169_Nuclei Isolation for Single Cell ATAC Sequencing’. Single-cell ATAC libraries were prepared according to the manufacturer’s user guide ‘Chromium Next GEM Single Cell ATAC Reagent Kits v1.1’. All libraries were sequenced on Illumina HiSeqX 150 bp PE with 1% PhiX.

### Microbiome analysis

Faecal samples were stored immediately at -80°C, and bacterial DNA was extracted after centrifugation and supernatant removal according to previous reports with a minor modification [94]. The V4 variable region was PCR-amplified with universal primer pairs (515F–806R) and the PCR product was sequenced on an Illumina Miseq. Libraries were prepared following the manufacturer’s protocol (Takara Bio, Inc.). A sample library with 20% denatured PhiX spike-in was sequenced by Miseq using a 500 cycles kit to generate 2 × 250 bp paired-end reads.

Data were processed using the DADA2 package (version 1.20.0) in R (version 4.1.1) following the standard pipeline [95]. Unless otherwise specified, default parameters were employed throughout. Briefly, reads were filtered and trimmed based on quality scores, using *filterAndTrim()*. Error rates were learned independently for each direction with *learnErrors()*. Amplicon sequence variants (ASVs) were inferred using *dada()*. Forward and reverse reads were merged with *mergePairs()* to reconstruct the full denoised sequences. Chimeric sequences were identified and removed using the consensus method via *removeBimeraDenovo()*. The final ASV table was constructed via *makeSequenceTable()*. ASVs were taxonomically classified using the naïve Bayesian classifier implemented in DADA2 (*assignTaxonomy()*) against the SILVA v138.1 reference database. Species-level classification was performed, where possible, using *addSpecies()*.

### Metabolome analysis

We followed the hydrophilic metabolites extraction and measurement methods as previously described with minor modifications [96]. In brief, methanol:water:chloroform was added in a ratio of 5:5:2 to 10 μL of plasma; 5 mg of faecal samples; and 5 mg of lung, mesentery, terminal ilium part of the small intestine, large intestine, and back skin. Tissue samples were homogenized using the Shake Master NEO (Bio Medical Science) with 3 mm zirconia beads at 1,500 rpm for 5 min. The extracted solutions from all steps were evaporated dry using a vacuum evaporator and lyophilized using a freeze dryer. Dried extracts were derivatized with methoxyamine hydrochloride (Merck) and N-methyl-N-trimethylsilyl-trifluoroacetamide (MSTFA, GL Science) before GC/MS/MS analysis, which was performed using a gas chromatography–tandem mass spectrometry (GCMS) platform on a Shimadzu GCMS-TQ8040 triple quadrupole mass spectrometer (Shimadzu) with a capillary column (BPX5) (SGE Analytical Science). The GCMS analysis and GC/MS/MS analysis were as previously described [96]. Normalised concentrations were compared by an unpaired two-tailed t-test. Multiple testing correction was performed with the BH method.

### Immunofluorescence

#### Labelling of alveolar macrophages (AM)

Mice were sacrificed by cervical dislocation, and the lungs were perfused with 5 mL of PBS followed by 5 mL of ice-cold PFA (4%), at a perfusion rate of 1 mL/min. Subsequently, whole lungs were collected and fixed in PFA (4%) for 24 hours. The lungs were then washed with PBS for 3 hours to remove the PFA before the lobes were embedded in agarose (2%) at 45° C. Embedded lungs were sliced (100 µm thick) using a vibratome (Leica, VT1200S). Lung slices were permeabilized in Permeabilization/Blocking Buffer (Ce3D™ Tissue Clearing Kit) for 24 hours at 4° C, followed by a 2-hour wash with Wash Buffer (Ce3D™ Tissue Clearing Kit) at room temperature. Before staining, lung tissue sections were treated with bleaching buffer (100 mM NaHCO_3_, 3% H_2_O_2_) for 30 mins at room temperature to minimise autofluorescence. For labelling, lung tissue slices were incubated at room temperature for 24 hours with AF647-Lectin (1:200; to label ECs), PE-anti-SiglecF (1:200; to label alveolar macrophages), and DAPI (1:1000; nuclear marker). Following incubation, the slices were washed with Wash Buffer for 2 hours at room temperature. Tissue Clearing Solution was then applied for 1 hour, after which the sections were immediately imaged using a Zeiss confocal microscope.

#### Quantification of AM numbers

Quantification of numbers and 3D rendering of alveolar macrophages in confocal images was conducted using Imaris software functions, Spots function, and Surface function. For each mouse, 10 complete alveolar macrophages were randomly selected, and their volume and sphericity was quantified.

#### SA-β-Gal staining

Lung tissue slices were incubated with the CellEvent Senescent Green probe (1:1000) at 37° C for 5 hours, followed by a 1-hour wash with Wash Buffer at room temperature. To control for inherent autofluorescence of alveolar macrophages, signals attained from mouse lung tissue slices incubated in a buffer solution at 37° C for 5 hours in the absence of CellEvent Senescent Green probe were subtracted from the experimental samples. The lung slices were then fixed with 4% PFA for 10 min at RT and alveolar macrophages were labelled with PE-anti-SiglecF, as detailed above.

#### Quantification of SA-β-Gal

Control images from young and aged mice, acquired from samples prepared in the absence of the CellEvent Senescent Green probe buffer solution, were used to set the image acquisition threshold. Fluorescent intensity of SA-β-Gal was measured in complete alveolar macrophages (randomly selected), subjected to 3D surface rendering (Parameter = 0.2).

### Single-cell data analysis

#### scRNA-seq data processing

As we included short fragments during library preparation, some Read 2 sequences did not contain genomic inserts suitable for mapping. To address this, only the original Read 1 sequences were used for downstream data processing. Specifically, using seqtk, the first 110 bp of each original Read 1 FASTQ entry was extracted and designated as R1, while the reverse complement of the last 80 bp of the same Read 1 was generated and used as R2. These modified R1 and R2 sequences were then used as input for the Cell Ranger (version 6.0.0) pipeline.

For each scRNA-seq library (n=141), gene-barcode matrices were generated using *cellranger count* with the reference genome refdata-gex-mm10-2020-A, obtained from 10x Genomics resources. Ambient RNA were then removed by CellBender (version 0.2.0) [97] and doublets were removed by Scrublet (version 0.2.3) [98]. Quality control was performed using the Seurat Package (version 4.0.4) [99]. Count matrices for all libraries were combined (384,381 cells x 27,755 genes) and filtered for cells with >1000 UMI, >500 genes, <20% mitochondrial reads, and genes expressed in >200 cells, resulting in 180628 cells x 27755 genes for downstream analysis.

We performed batch correction and integration by using the SCVI package (version 1.2.1 [100]) with Scanpy (version 1.10.4 [101], python 3.11.0) on the raw counts. The model was set up to perform integration per library and we trained the model with n_layers=3 and n_latent=50, with other settings default. Nearest neighbours were calculated using the SCVI reduction, and the UMAP was determined (min_dist=0.3, default settings). To generate clusters with unsupervised clustering, Leiden clustering was performed (flavor=’igraph’, n_iterations=10, resolution=1.3, leidenalg version: 0.10.2).

#### scRNA-seq cell annotation

scRNA-seq data were annotated by visualising marker expression across Leiden clusters by using cell type <u>marker genes</u>. An initial set of annotations was collected by grouping T cells, B cells, intestinal epithelial cells, myeloid cells, neural crest cells, hair follicle cells, and gland cells. These grouped compartments were subannotated by isolating the set of cells and performing Leiden clustering and a new UMAP embedding. Subcluster annotations were made to generate a level 2 annotation depth. Lastly, fibroblasts, endothelial cells, intestinal epithelial cells (not stem cells), and secreting cells were separated by tissue origin as we observed significant tissue differences. Clusters deemed low quality were removed. Lastly, we generated UMAPs for alveolar macrophages and other epithelial cells for further analysis.

#### scATAC-seq data processing

FASTQ files of scATAC-seq experiments were processed using Cell Ranger ATAC (Version 2.0.0) (Intestine samples: version 2.1.0, reference mm10-2020-A-2.0.0. We utilised the ArchR toolkit due to the size of the cell per peak (version 1.0.2) [85]. TThe ArchR toolkit generates arrow files per library from the fragment files to save data as a system file instead of per memory. Chromatin accessibility is binned per 500 bp for initial processing and clustering, prior to peak calling. The integrated doublet finder method of ArchR was applied to remove doublets. Additional filter parameters were >10 TSS enrichment and > 3500 fragments per cell (nfrag), and we applied a whitelist of Cell Ranger cell calling (filtered barcode list). Dimension reduction and clustering were performed using the ArchR built in tools using the binned matrix. In brief, we performed “additerativeLSI” on the tilematrix (iterations: 4, dimsToUse 1:80, varFeatures: 25000) to generate reduced dimensions. Clusters and UMAP embedding were generated with addClusters (resolution 2, max clusters 100) and addUMAP (metric: cosine).

#### scATAC-seq cell annotation

We performed label transfer with addGeneIntegrationMatrix per tissue using geneScoreMatrix. Label transfer of V3, per tissue (other than intestines) cell types (fibroblasts, endothelial cells) were merged. Profilerating and non-annotated cells were removed, then label transfer was performed tissue by tissue. We conducted consensus-based relabelling in which clusters comprising over 80% of a cell type were labelled to that cell type. Clusters between 60% and 80% also underwent this process, except for cluster 60 (pre B cell, immature B cell); keratinocytes; and hair follicle cells. Annotations were further cleaned up by adding erythrocytes and renaming clusters with <60% consensus. Finally, fibroblasts and endothelial cells were separated by tissue. We removed a cluster containing mixed accessibility markers of T and B cells (Cd8a+ and Cd19+) as we were not able to properly annotate these cells.

#### scATAC-seq peak set and motif annotations

We performed group coverage of annotated cell types with the “addGroupCoverages” function. A reproducible peak set was generated with MACS2 using the “addReproduciblePeakSet” function (peaksPerCell = 1000, maxPeaks = 1000000, other settings default). Motif annotations were generated using the “cisbp” motif set and the “addMotifAnnotations” function. We opted not to use the ENCODE blacklist regions as we observed regions of genes of interest such as Mid1 overlap with the blacklist regions, as well as age-dependent changes. We did not observe global read in blacklist ratio changes.

#### Integrative analysis

To integrate the scRNA-seq data with the scATAC-seq data, we gathered the scATAC-seq genescore matrix stored in the ArchR arrow files. In order to obtain the integrated scRNA-seq and scATAC-seq combined UMAP embedding, CCA anchor transfer was used with the Seurat package, for 5000 variable RNA genes. The genescore matrix generated with ArchR was used as a proxy for scATAC-seq.

#### Cell proportion analysis

Cell type proportion was calculated by the number of cells of each cell type divided by the total number of cells, for each tissue in each mouse separately. We compared the proportion for each cell type in each scRNAseq tissue between (1) old SPF / young SPF and (1) old GF / young GF. The effect size was evaluated by the logarithm ratio of fold change (log2FC), and the p-value was determined with a GLM model using a quasibinomial family. Multiple testing was corrected with a BH post-hoc.

#### Differential expression and directionality

For each cell type (in all tissues and all mouse replicates), we performed exploratory differential expression analysis (*Seurat::FindMarkers()*) for the following pairwise comparisons: (1) old SPF / young SPF, (2) old GF / young GF, (3) old GF / old SPF, and (4) young GF / young SPF. We used the Wilcoxon rank-sum test with a BH post-hoc correction. We used an alpha of 0.05 to determine significance. The Spearman correlation (*cor(method = "spearman")*) of fold change for all genes between (1) old SPF / young SPF and (2) old GF / young GF was calculated separately for each cell type.

To group DEGs as directional genes, we evaluated whether genes had significant similar directionality to compare corresponding cell types between natural and sterile ageing. If increased only in natural ageing, genes are called extrinsic; if genes increase only in sterile ageing, they are called sterile-only. If a gene changes significantly in both comparisons, it is termed intrinsic, and if the directionality reverses, the gene is called reversed. Additionally, when analysing generic ageing genes in the directional categories, we added a minimum cell type cut-off of more than 3 cell types and a LFC threshold of 1.25. These genes are then separated by whether they are upregulated or downregulated.

In order to score similarity between cell types, we compared the same directionality without a LFC cut-off per cell type. This analysis used genes that change in a similar direction for natural ageing and compared these genesets to those from sterile ageing, the effect of old GF, and the effect of young GF. We then determined similar versus opposite direction. For similarity scoring, we employed the Sørenson similarity index. We chose the Sørenson method over the Jaccard method to emphasize intersecting genes, as gene set sizes may behave differently due to the sizes of differential genesets, which are partly affected by cell type population sizes. To cluster cell types and similarity scores, we applied hierarchical clustering and k-means clustering using the complexheatmap package (version 2.12.1).

To validate potential candidate genes (Figure 4G-H, S4G-I, S5E), DEG analysis was performed at pseudobulk resolution using the DESeq2 package (version 1.36.0). Raw counts were aggregated using the *AggregateExpression()* in Seurat across mice replicates with tissues combined and per tissue comparisons. DEG analysis was performed using standard DESeq2 pipeline with default settings, with a genefilter of a minimum of 10 counts. The experiment was set up using *∼ model_age* as design included all conditions per cell type. For comparison between alveolar senescent states, a subset was made of the specific comparison prior to DEG analysis. The “*LFCshrink*” function was performed with the “normal” parameter to shrink LFC values.

#### Differential accessibility

To determine accessibility changes across conditions in different cell types, we performed a differential peak analysis for each cell type. Next, using the TxDb.Mmusculus.UCSC.mm10 reference (versions: ensGene_3.4.0, knownGene_3.10.0), we annotated the location type in the genome for each differential peak (Promoter, exon, intron, intergene). The promoter region type is defined as 200 bp upstream with a max gap between peak and promoter region of 300 bp upstream or downstream. The closest located gene per differential peak was determined by using the “*nearest()”* function.

#### Pathway activity

Geneset enrichment analysis of gene ontology biological processes was performed using the AUcell package (version v1.16.0) [46], where the area under the curve was determined per geneset across cells. Geneset scores for each cell type between conditions were further investigated by determining whether there is a significant difference using the Wilcoxon rank-sum test (alpha = 0.05) with multiple test correction by the BH method. To identify the most changing pathways, we ranked pathways by significance occurrence in cell types per pathway. To calculate senescence score, we applied the senmayo geneset [49].

Sets of genes, such as ageing genes, were characterised through enrichment of gene ontology gene sets (biological processes) by using the clusterprofiler package [102]. Significant genesets were calculated according to either upregulated genes or downregulated genes between conditions, or per cluster (p value < 0.05, q value < 0.05, p.adjust value < 0.01). Senescence was scored using the SenMayo gene set, and we used the list of genes curated for mice.

#### Alveolar macrophage and senescence analysis

To identify putative senescent cells in the alveolar macrophage population, we used a quantile cut-off as we treated young cells as non-senescent. Alveolar macrophages were isolated and senescent cells were annotated based on an alpha = 0.05 cut-off where 95% of young cells were “non-senescent” to account for false positives. The remaining 5% of senescent-scoring young cells were not labelled senescent. Old SPF and old GF samples were analysed separately.

#### Regulon activity and motif scoring

Regulon scores for each TF based on scRNA-seq gene expression per cell was calculated using the pySCENIC package (version 0.11.2) [46], and TF motif activity score based on scATAC-seq motif enrichment per cell was calculated using the chromVAR tool built in ArchR [45]. Motif annotations were used with the previously defined “cispb” motif set. Deviations were determined per TF per cell and scores were scaled (Z-score). Transcription factor activity change was further investigated to identify significant changes by using Wilcoxon rank-sum test (alpha = 0.05) with multiple test correction via the BH method. Transcription factors of interest were determined by how often the transcription factor was significant in the various cell types, comparing the conditions. These transcription factors of interest were visualised using pseudobulk values (mean per cell type per mouse replicate).

#### Immune repertoire analysis

To calculate clonal types of B and T cell receptors within 5’ scRNA-seq data, we used the Trust4 package per library [62]. We filtered for cells that were assigned T cell in both Trust4 and manual cell annotation and performed the same for B cells. As only a handful of cells were typed 100%, we used putative clonality using the VDJC (chain 1) genes for B cells, including chain 1 genes and the annotated CDRs. We applied clonality frequency as a fraction of the total B cell population, "> 1%", "0.1% < 1%", "0.05% < 0.1%","< 0.05%" by "Hyper Expanded", "Expanded","Minimal","Rare", respectively [103]. In the frequency analysis of B cell clonality, B cells without clonal typing as defined by us were removed.

#### Imputed single cell metabolomics

Quantitative System Metabolism (QSM) pipeline was outsourced to Doppelganger Biosystem GmbH [51].

#### Utilised publicly available single cell data

Analysis performed using publicly available data from: https://figshare.com/projects/Molecular_hallmarks_of_heterochronic_parabiosis_at_single_cell_resolution/127628 and https://cellxgene.cziscience.com/collections/0b9d8a04-bb9d-44da-aa27-705bb65b54eb. We performed the standard Seurat pipeline and isolated alveolar macrophages by marker expression (Siglecf+, Marco+).

## Resource Availability

### Lead contact

Requests for further information and resources should be directed to and will be fulfilled by the lead contact, Aki Minoda.

### Materials availability

This study did not generate unique new reagents.

## Acknowledgements

We are grateful to the animal facility of the RIKEN Yokohama campus for keeping mice germ-free for this study as well as Shunsuke Ishii and Piero Carninci for making this project possible as part of the RIKEN Aging Project. We also thank Natsuko Otaki, Takuma Misawa, and Masato Asaoka for guidance on cell type annotation. This work was supported by the RIKEN Aging Project and JSPS KAKENHI (20H03692) to A.M. S.N. is funded by a Wellcome Trust Investigator Award (221699/Z/20/Z).

## Author contributions

A.M. conceived the study and supervised the project. A.M. and T.W.T. designed the experiments and coordinated the study.

H.Y., S.K., N.H., P.J., T.K., S.M., M.M., Y.M., Y.N., M.A., N.T., K.T., M.V.B., N.S-T. collected and processed tissues.

H.Y. and S.K. generated single-cell data.

T.W.T, J.C, U.T. and T.H. performed cell type annotation.

W.N.T., J.C., and T.W.T performed data curation and analyses.

H.T.W. and S.N. designed and performed AM senescence imaging experiments.

N.R. performed ELISA experiments.

A.A. performed SCAFE and scATAC data analyses.

N.S-T., H.F., T.M., K.M. and H.O. contributed to study design.

C.C.H. contributed to data analyses.

A.M. and W.N.T. wrote the manuscript with inputs from J.C. and T.W.T.

## Declaration of generative AI and AI-assisted technologies in the writing process

While preparing this work, the author(s) used ChatGPT 4o and 5.1 (OpenAI version 5.1 2025) in order to improve manuscript readability. After using this tool/service, the author(s) reviewed and edited the content as needed and take full responsibility for the content of the published article.

