## Supplemental Figures for "Epigenomic and transcriptomic germ-free ageing atlas reveals sterile inflammation as an intrinsic ageing feature"

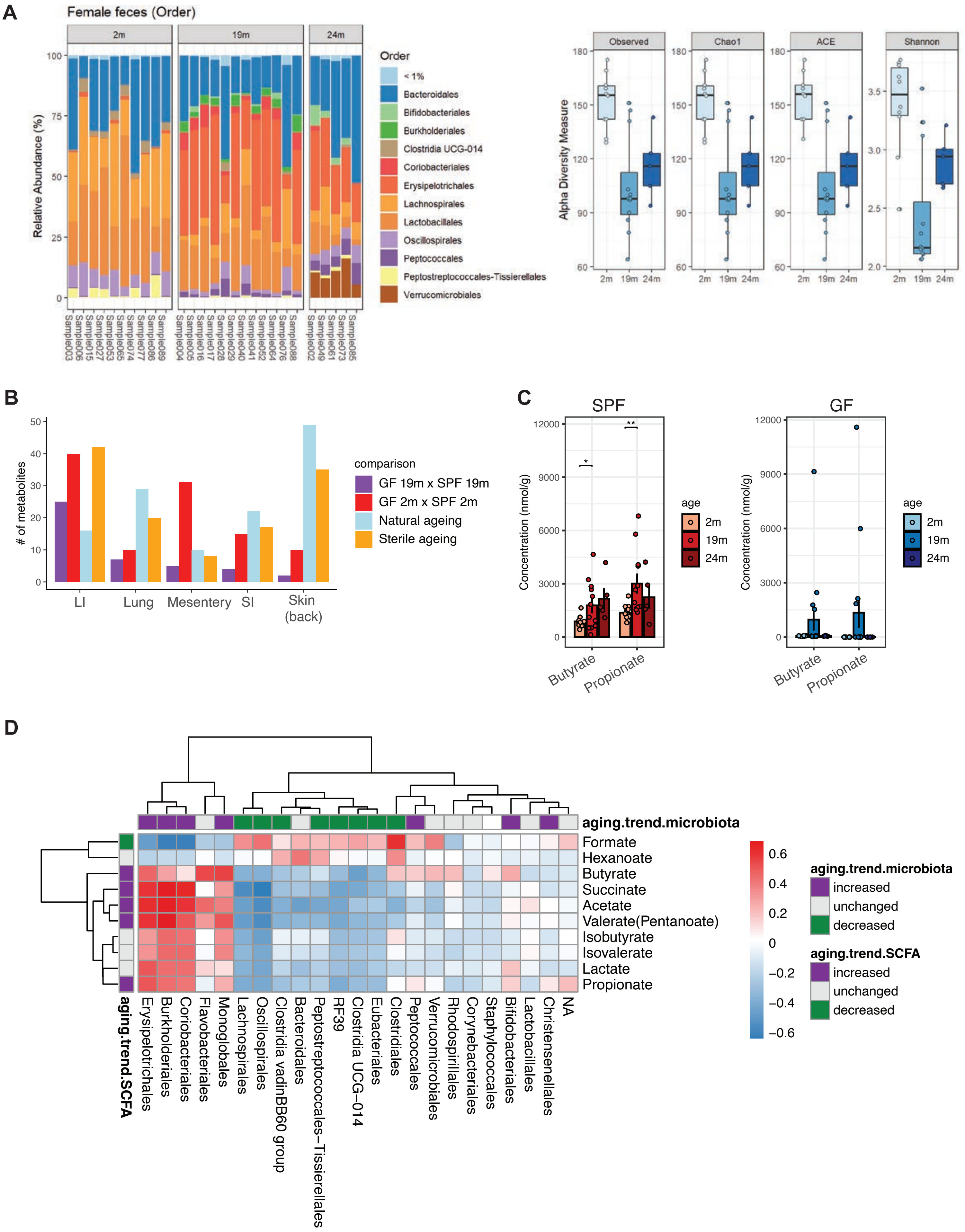


**Figure S1. Faecal samples analyses. Microbiome and metabolites changes with ageing . A.** Fractions of bacterial species from fecal samples across SPF mice (sample) at the order level shows dysbiosis (left). Boxplot of alpha diversity scores for shown methods for faecal samples separated by age shows decline in community complexity (right). **B.** Significant number of metabolites detected across tissues and conditions by count (ageing: 19m vs 2m). **C.** Targeted mass spectrometry of SCFAs on facecal samples show indicated SCFAs are produced in a microbiota-dependent manner (absent in GF; technical noise is observed in GF 19m) and increase with old age in SPF. **D.** Pearson corrleation to compare changes in microbiota to SCFAs.

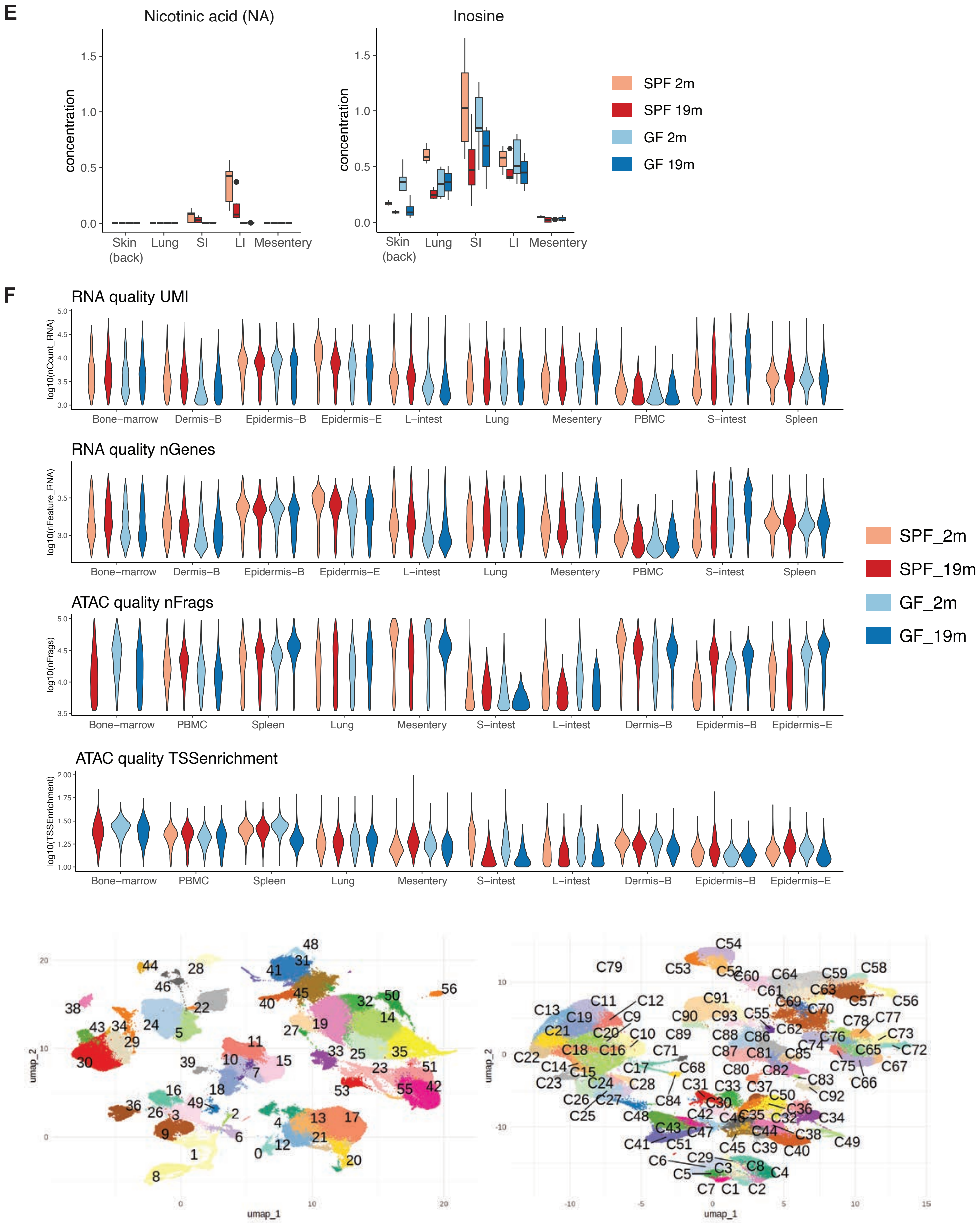


**Figure S1 (continued). Data quality and clusters. E.** Example metabolites changing with age and condition, separated by tissues and conditions. **F.** (Upper) Violin plots of quality parameters per cell per condition per tissue. UMI: RNA count of UMIs per cell. nGenes: number of detected genes per cell (RNA). nFrags: Count of fragments per cell in the scATAC-seq data. TSSenrichment: score of how many fragments should be detected in TSS regions relative to non-TSS regions. (Lower) scRNA-seq (left) and scATAC-seq (right) UMAP visualisation with unsupervised clustering labeled.

**G**

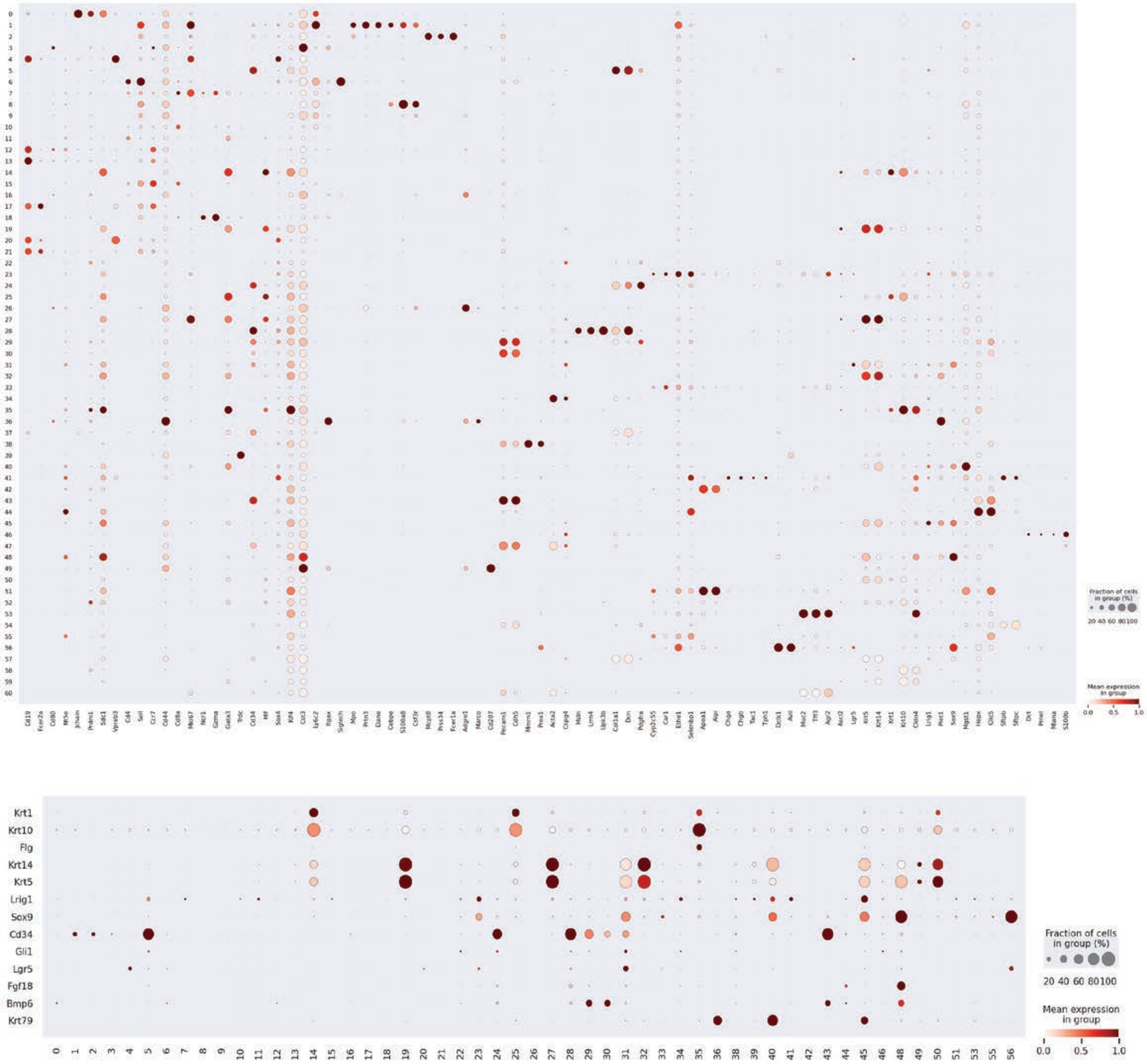


**Figure S1 (continued). G.** Dot plots of cell type marker expression across clusters. Upper panel shows for all and lower panel shows skin-specific markers.

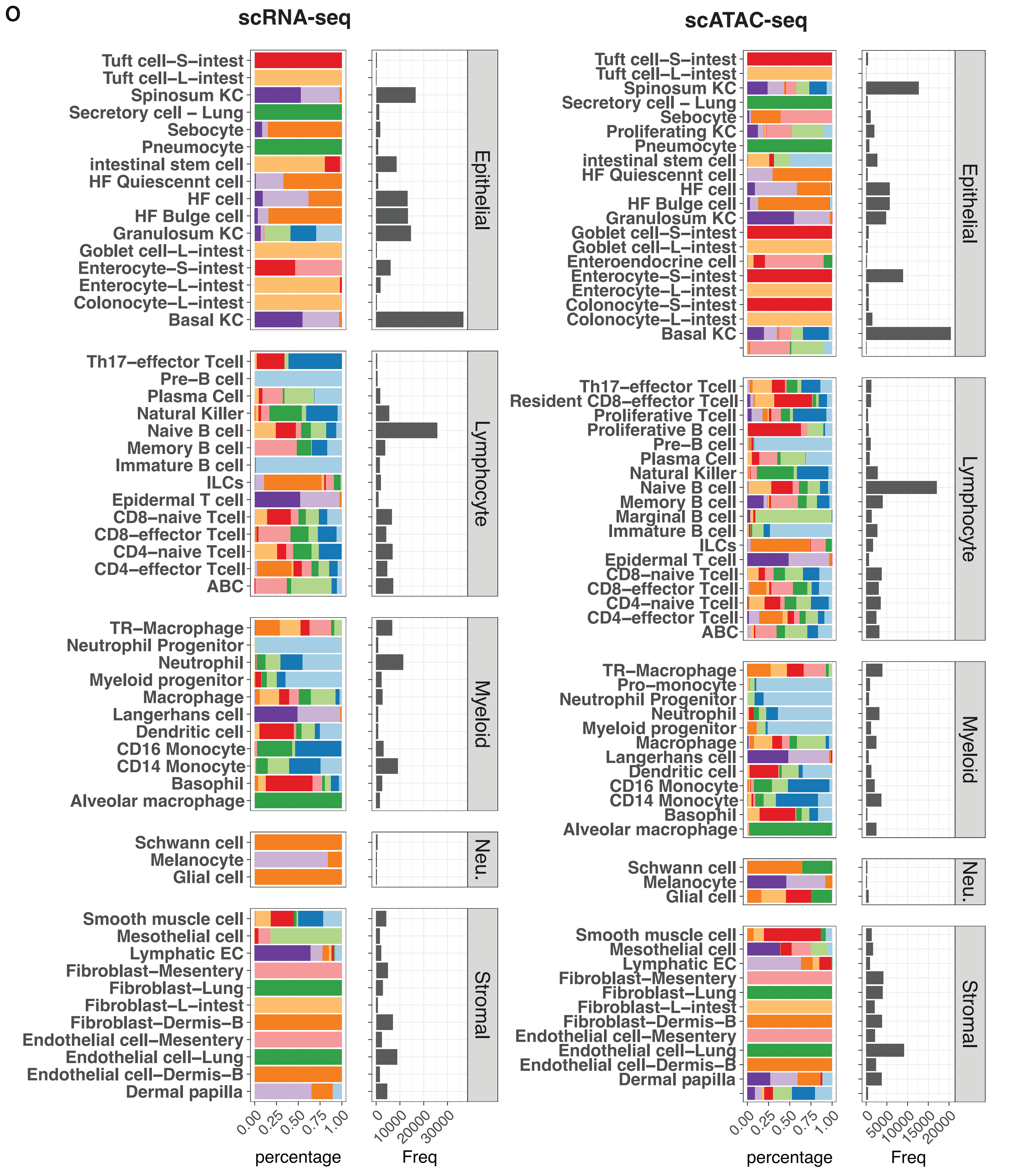


**Figure S1 (continued). Cell type proportions of the Mukin atlas. O.** Cell type proportions are shown for indicated compartments from scRNA-seq (left) and scATAC-seq data (right), origin of tissues are indicated by colours and total number of cells per cell type are shown in grey.

P

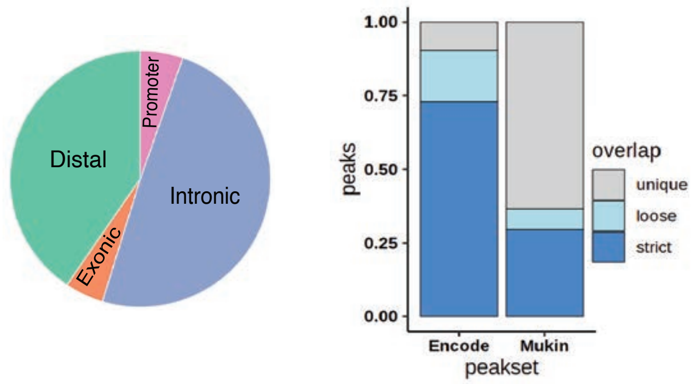


Q

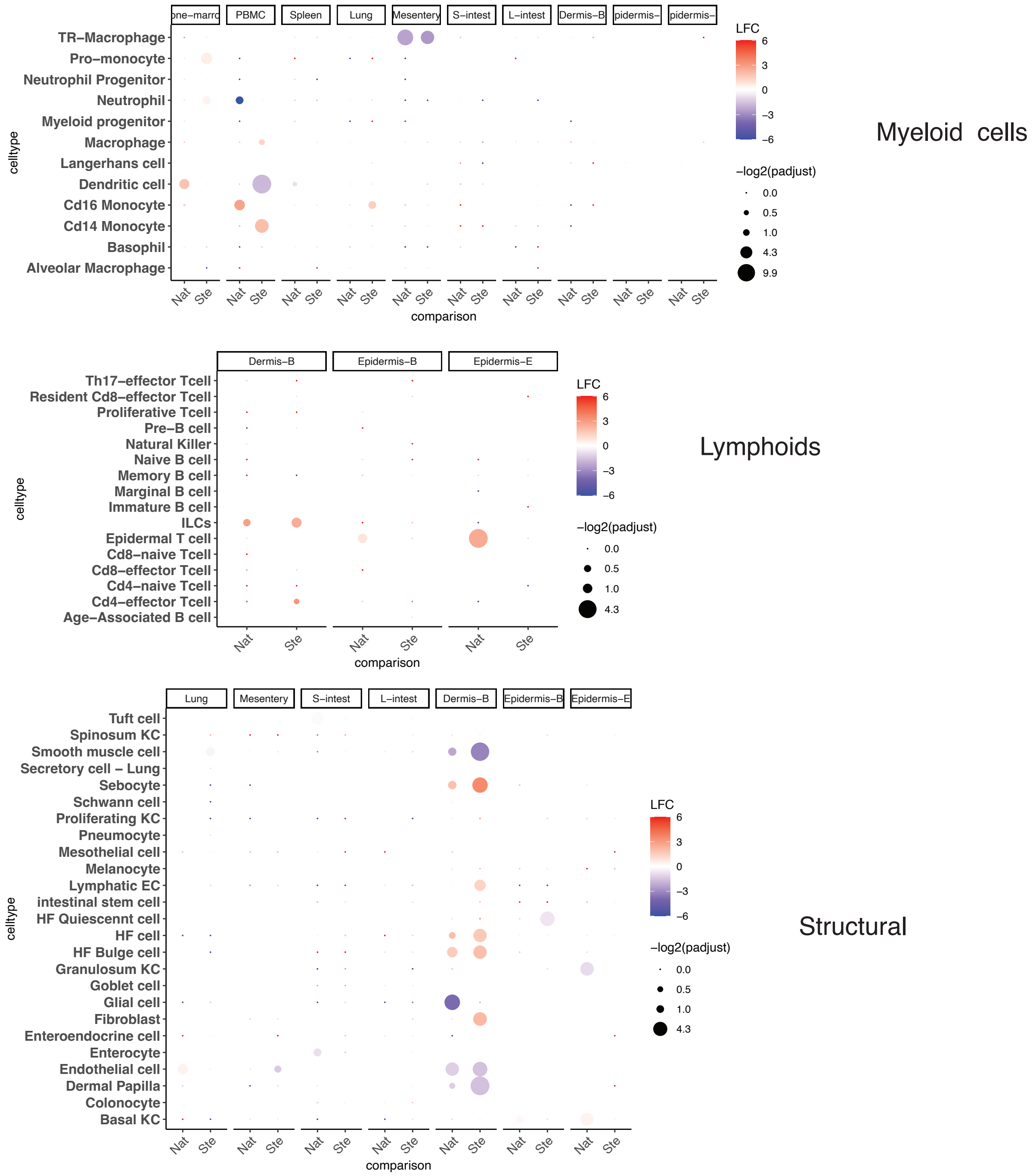


**Figure S1 (continued). Cell type proportion changes with ageing. P.** Pie chart distribution of annotated peaks in the scATAC-seq dataset (left) and comparison pf annotated peaks to ENCODE peak sets. **Q.** Cell type population changes across tissues by compartments as described. LogFCs are shown for natural (Nat) and sterile (Ste) ageing by color gradient. Dot sizes are scaled by significance (-log2(pvalue), pvalue: unpaired two-tailed t.test).

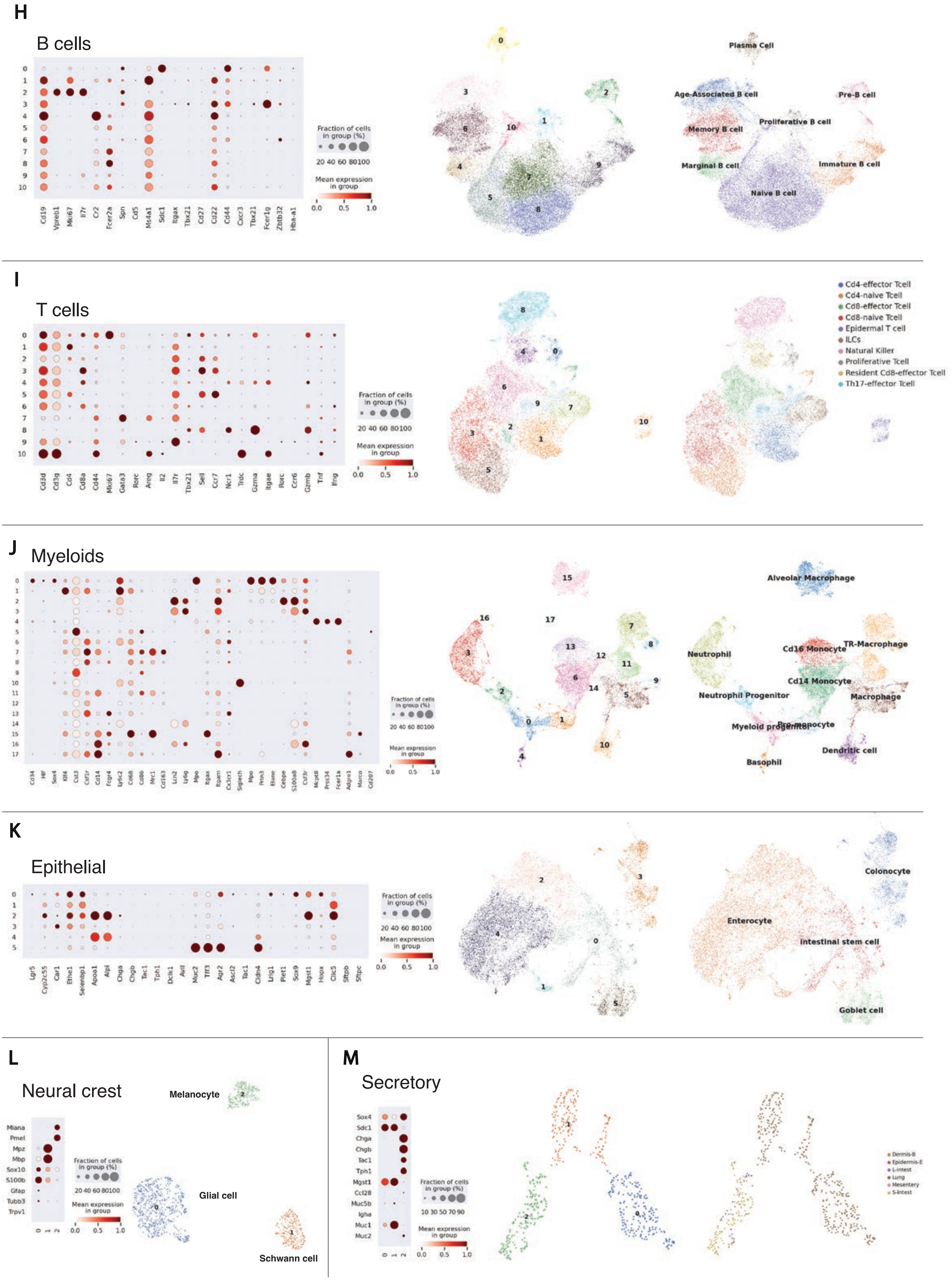


**Figure S1 (continued). Subcompartment clusters**. Unsupervised clustering (left), marker expression (middle), and cell annotations (right) of B cells (**H**), T cells (**I**), myeloids (**J**), epithelial (**K**), neural crest (**L**), and secretory (**M**).

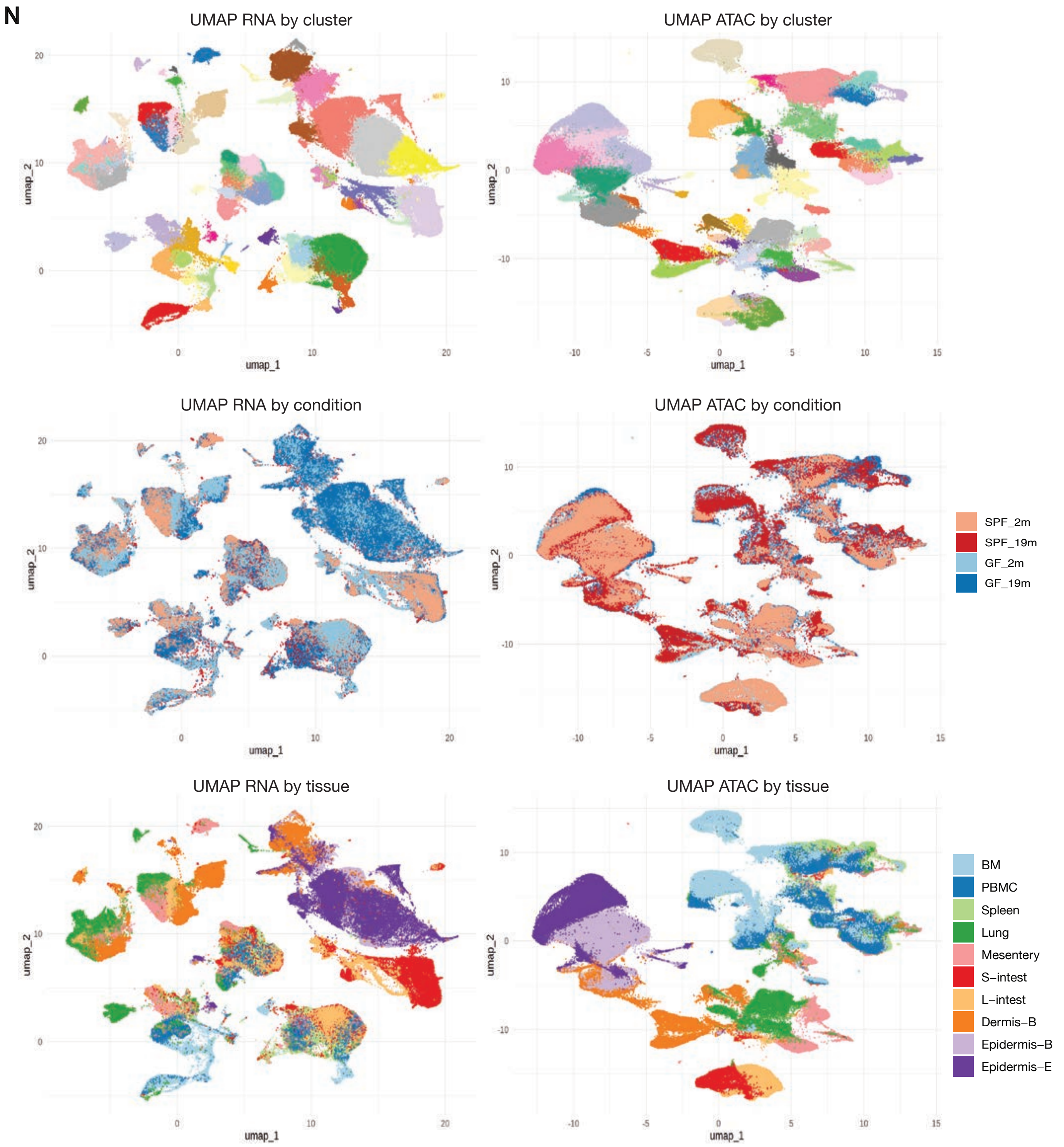


**Figure S1 (continued). N.** UMAP embeddings of clusters of scRNA-seq (right) and scATAC-seq (left) by clusters (first row), by conditions (second row), and tissues (third row).

**R**

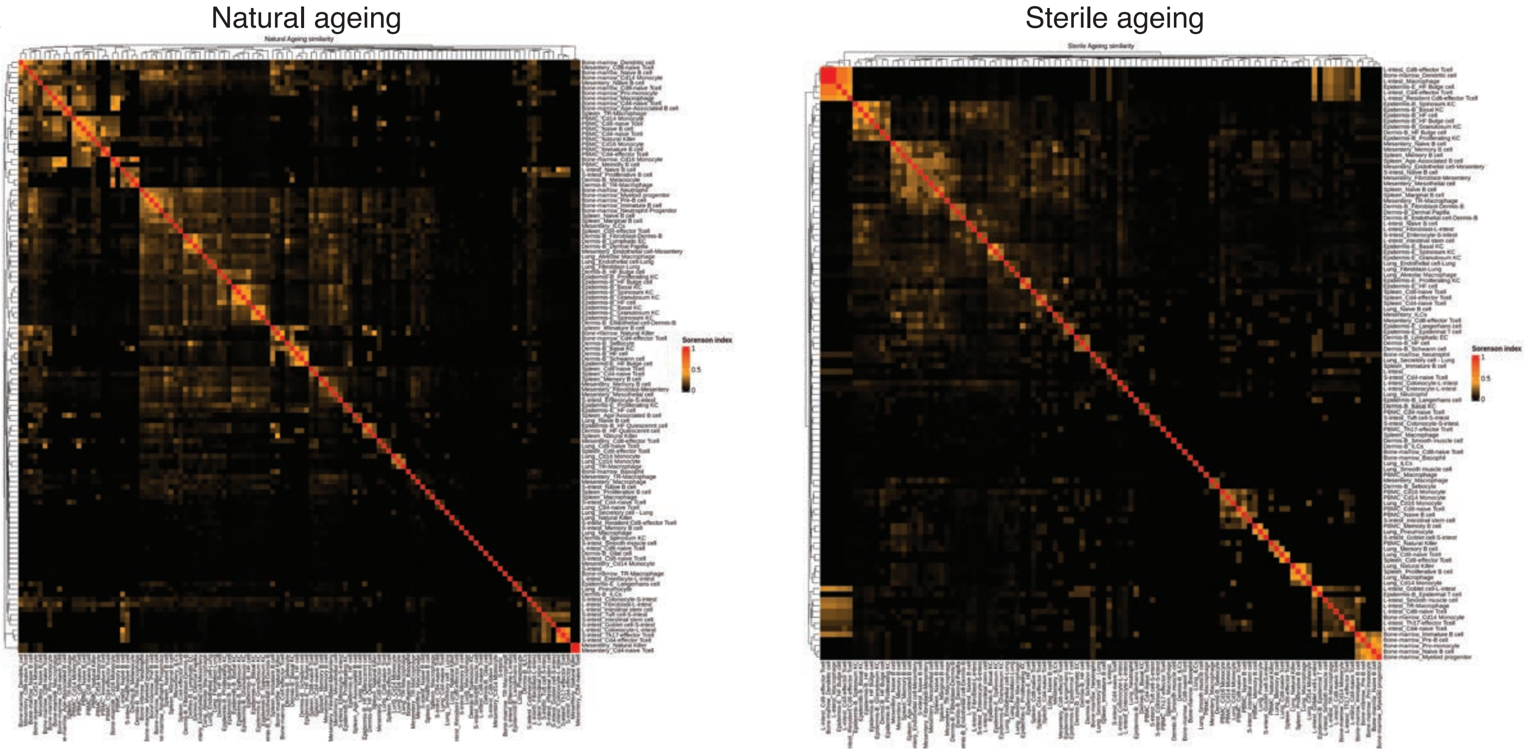


**S**

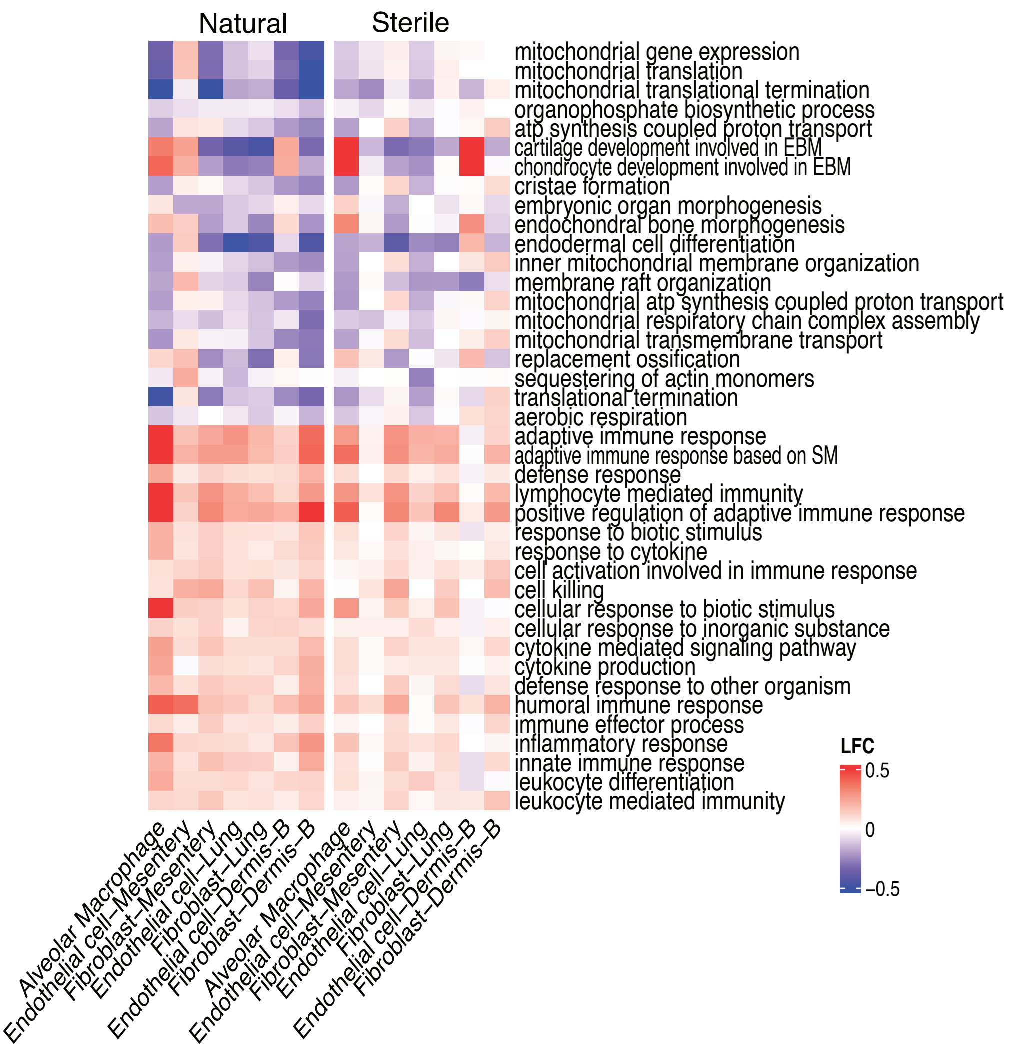


**Figure S1 (continued). Tissue ageing. R.** Sorenson similarity scores with DEGs sets between all cell types comparison for natural (left) and sterile (right) ageing. Heatmap is clustered by hierachical clustering. **S** Heatmaps showing LFC of genesets of interest across cell types in natural (left) and sterile (right) ageing.

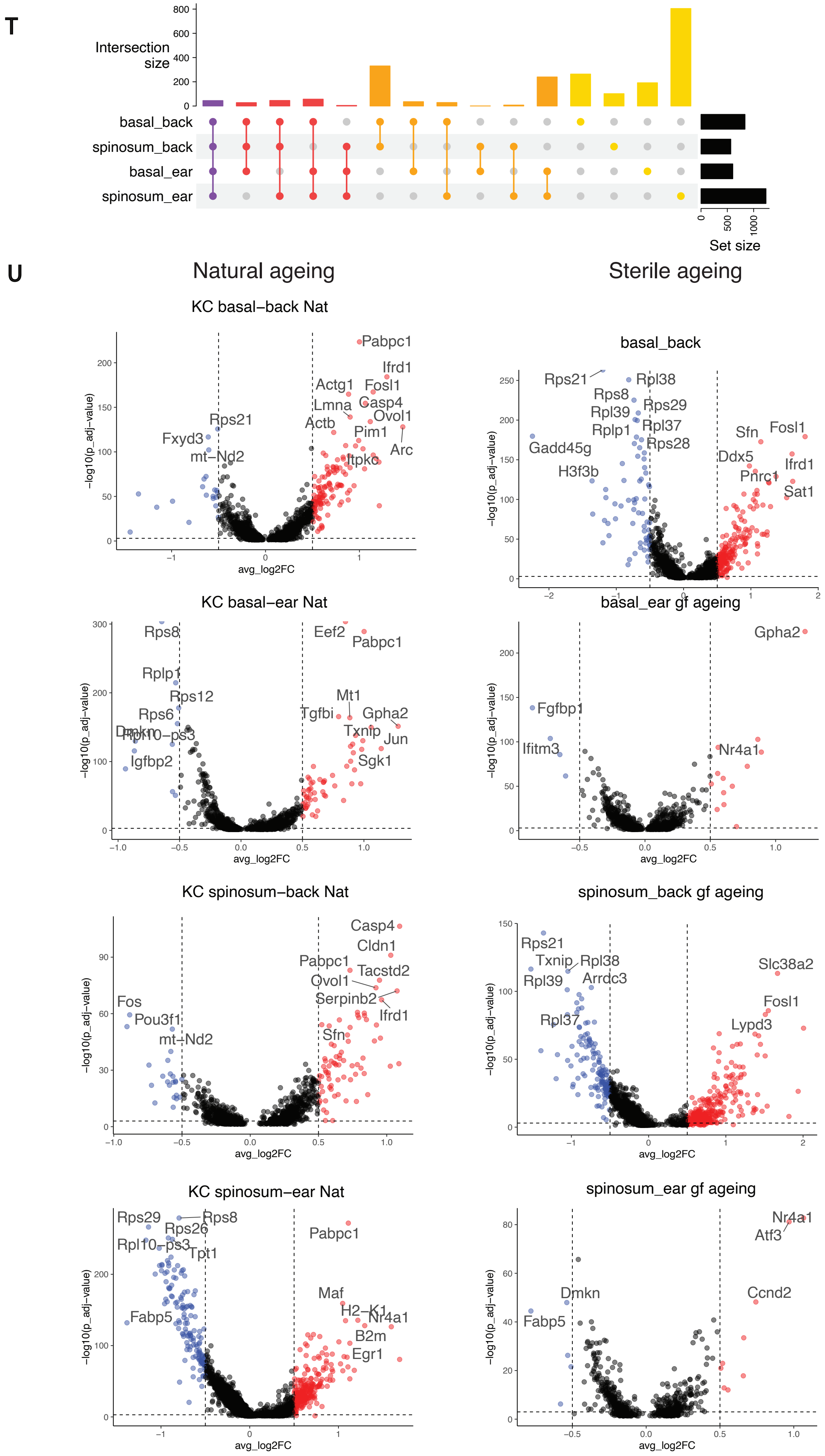


**Figure S1 (continued). Tissue ageing. T.** An upset plot of upregulated natural ageing genes in basal and spinosum KCs from back and ear epidermis. **U.** Volcano plots showing natural (upper) and sterile (lower) ageing DEGs for shown cell types. Upregulated (red) and downregulated genes are plotted by their respective log2 fold change and -log10 p.adjusted value (wilcoxon rank sum test). Various genes that were most signifi-

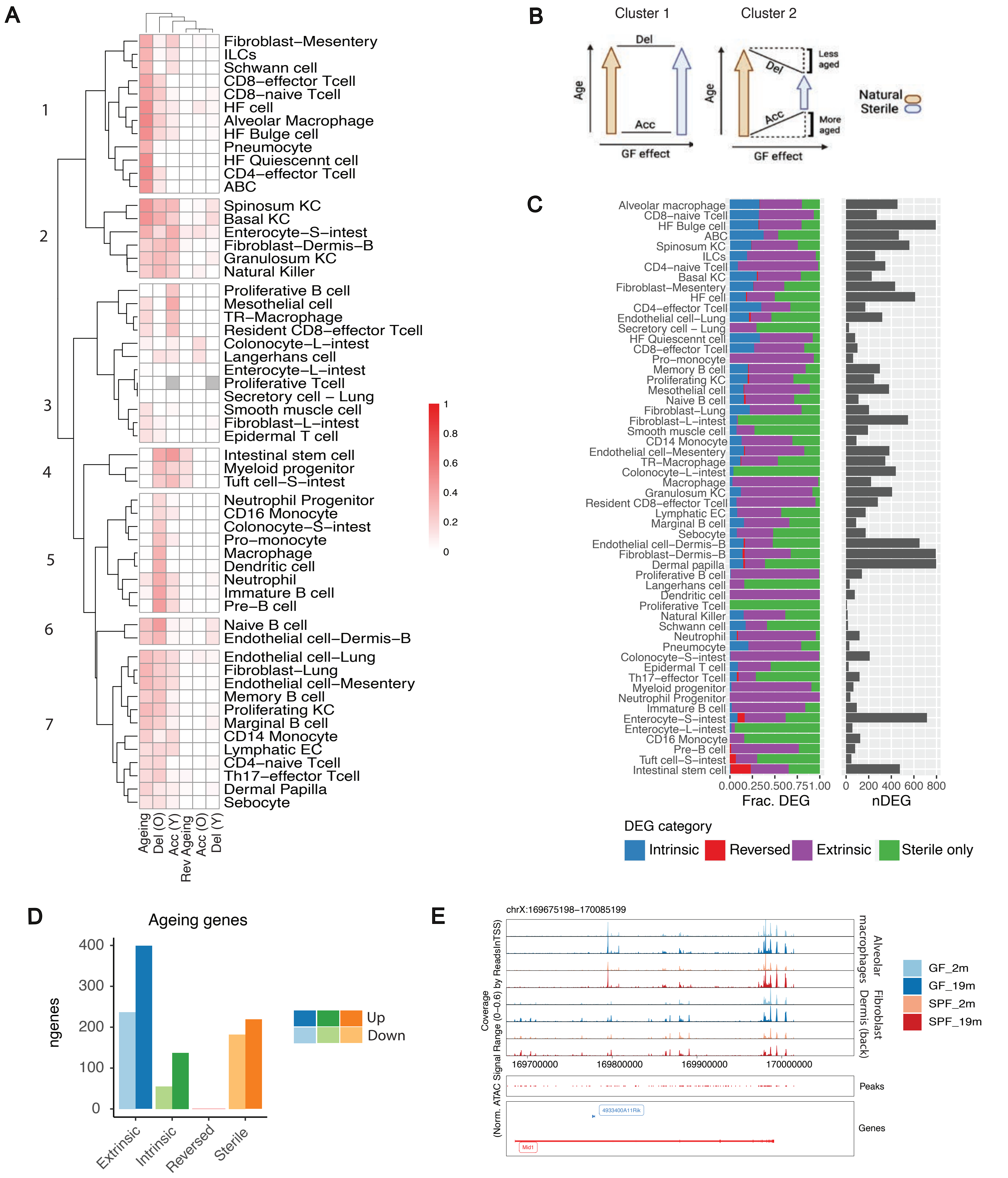


**Figure S2A-E. Transcriptomic profiling of microbiota effects during ageing. A.** “Ageing” column shows comparisons between natural and sterile ageing, reflecting the effects the GF condition has on ageing. A high similarity value reflects minimal effect and a low value reflects larger effect. High similarity score in ‘Del (O)’ column indicates whether ageing is delayed and is derived from similarity score between natural ageing DEGs to ‘old GF:old SPF’ DEGs. ‘Acc (Y)’ column shows the extent of accelerated ageing based on similarity scores between natural ageing DEGs to ‘young GF:young SPF’ DEGs. Sorenson scores were calculated separately for upregulated and downregulated gene sets. Hierarchical clustering and dendrogram clustering were subsequently performed for group cell types. **B.** A schematic of the relationship between natural ageing, sterile ageing and the (SPF old x GF old) and (SPF young x GF young) comparison as shown in Sorenson correlation. **C.** A directional analysis between LFC of natural ageing and sterile ageing. DEGs are binned into extrinsic (SPF only), sterile only (GF only), intrinsic (both SPF and GF) and reversed (opposite direction in SPF and GF ageing). Total number of DEGs per cell type are visualised in grey. **D.** Total number of ageing genes (differentially expressed in more than 3 cell types) per group of directional category. **E.** Genome browser showing normalised chromatin accessibility peaks for each condition in basal KCs at Caspase1/4/12 locus.

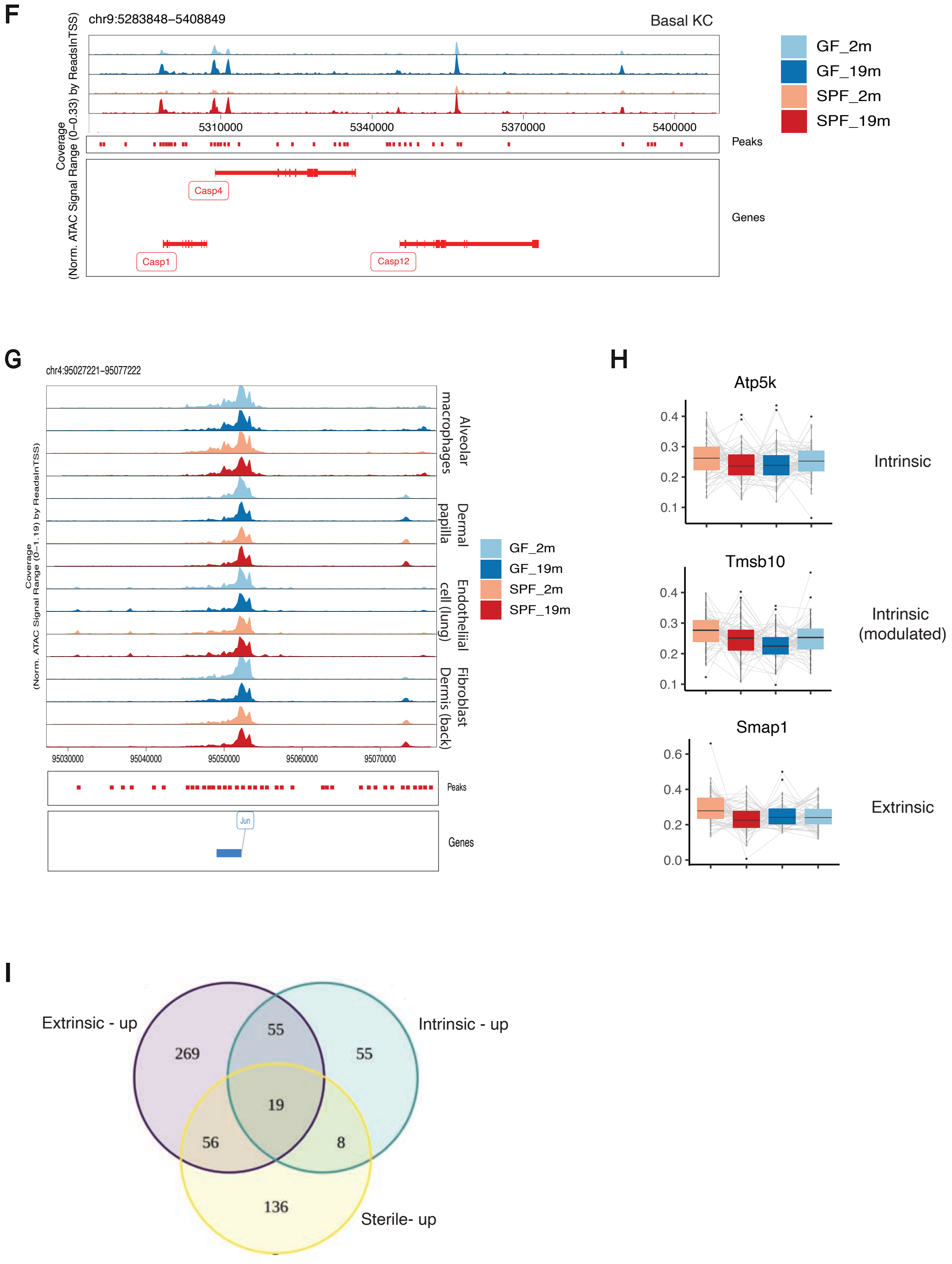


**Figure S2F-I. F.** Average expression per cell type visualised as a boxplot. **G-H.** Genome browser showing chromatin accessibility peaks (normalised accessibility peaks) for each condition for indicated cell types at Caspase genes (G) and Jun (H) loci. **I.** Venn diagram indicates number of upregulated ageing genes in each category, which shows overlapping genes between groups that are upregulated

**J**

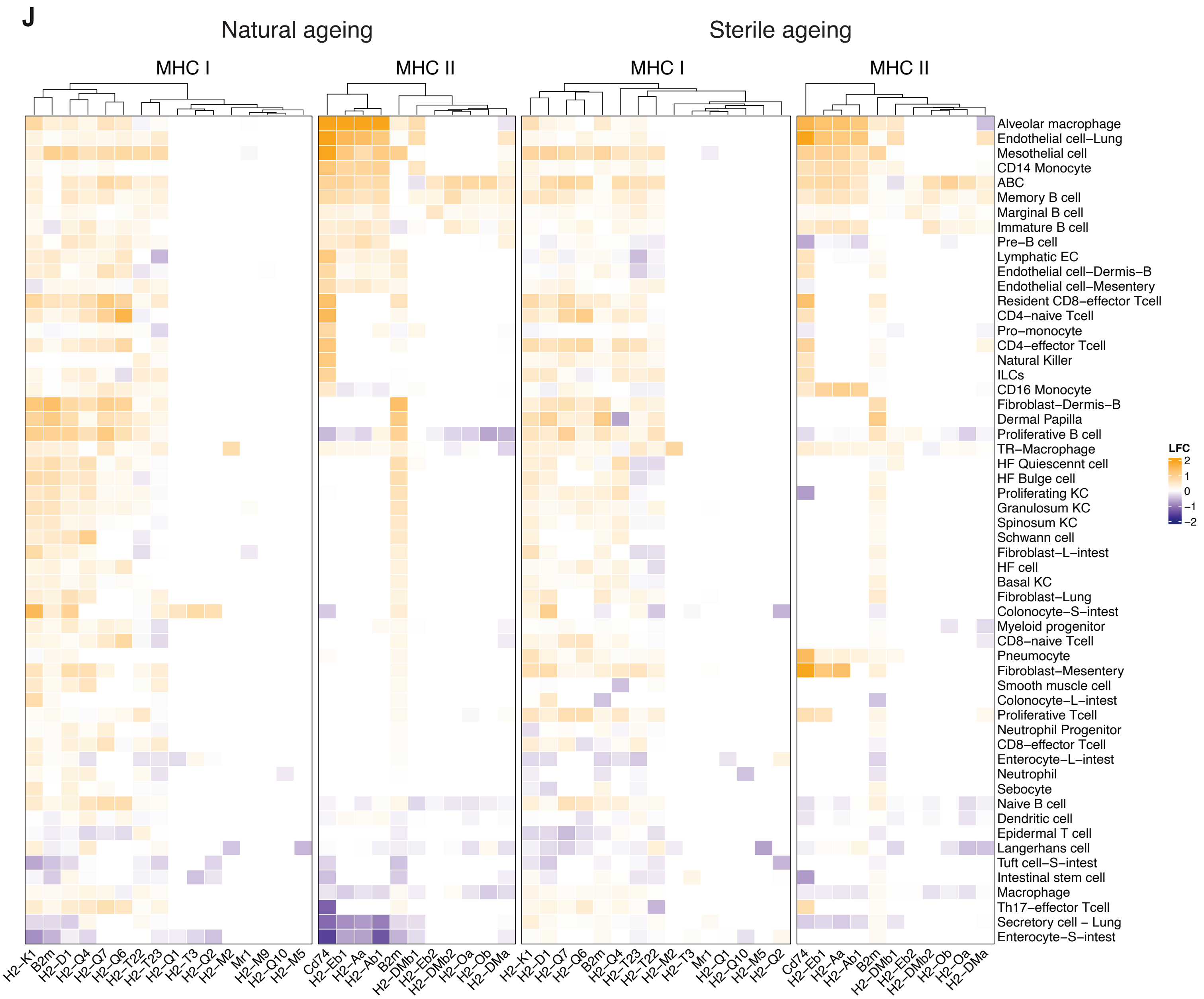


**K**

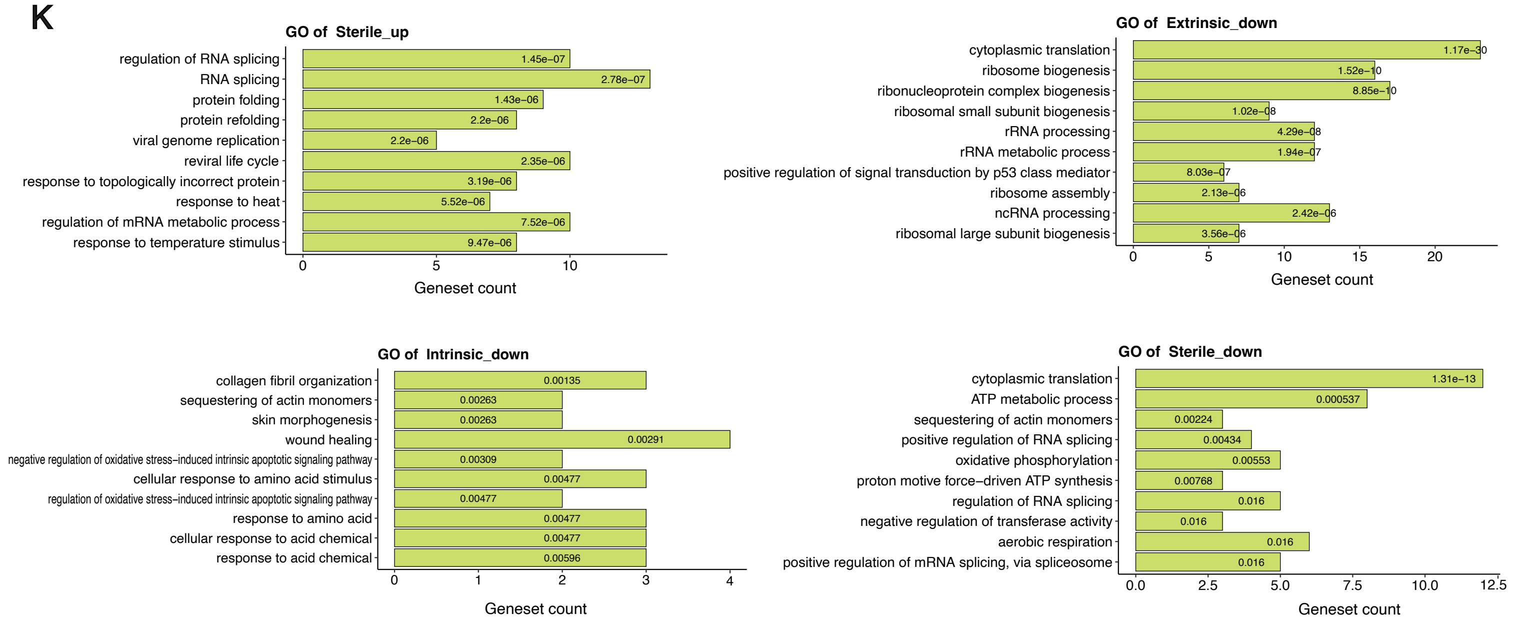


**Figure S2J-K. J.** Heatmap of LFC of all MHC I and MHC II genes for natural and sterile ageing, which shows they are dynamically deregulated during ageing. **K.** Top GO terms for each caterogry

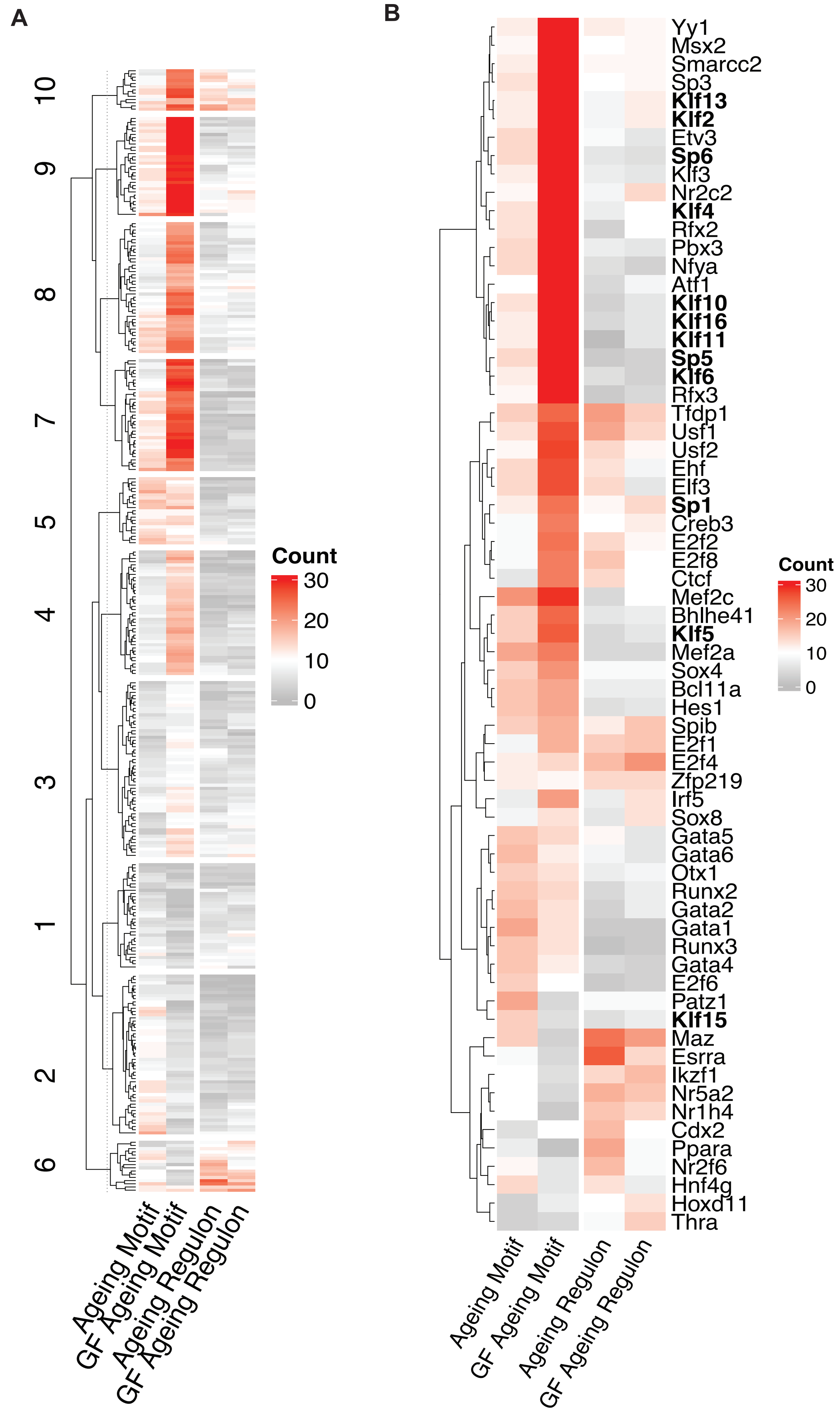


**Figure S3A-B. A.** Downreglated TF activities during natural and sterile ageing by means of motif and regulon activities. TFs are grouped and clustered based on hierarchical clustering and dendrogram clustering, respectively. Heatmap shows number of dysregulated cell types. Refer to Supplementary Table 8 for lists of TFs. **B.** Selected TFs from (A) are shown. Sp/Klf TFs are in bold.

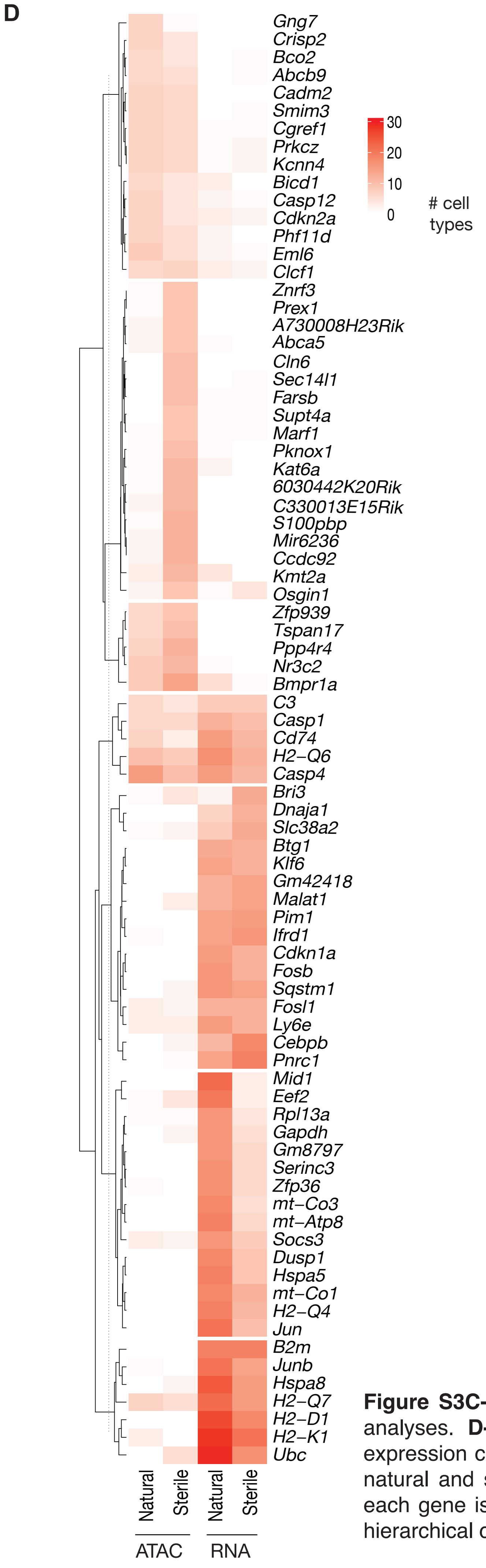


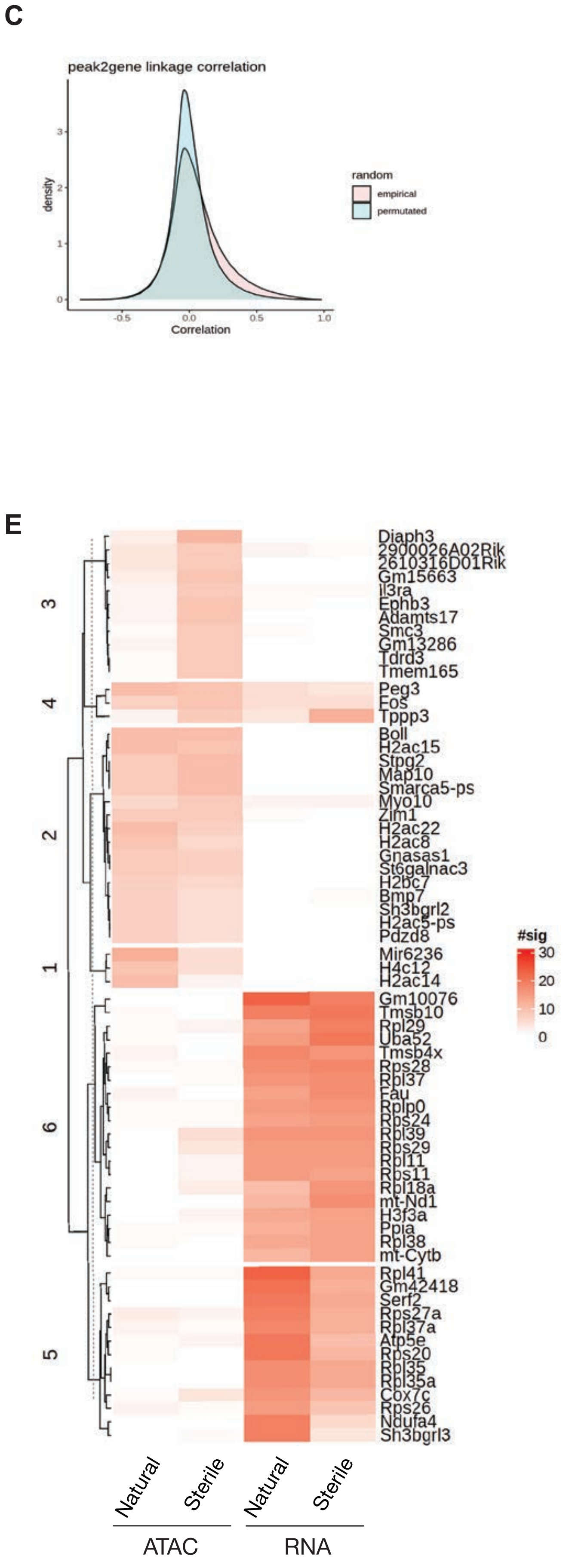


**Figure S3C-E. C.** Peaks were linked to the nearest gene for subsequent analyses. **D-E.** Correlation of chromatin accessibility (ATAC) and gene expression changes for genes that become open (D) and closed (E) during natural and sterile ageing. Heatmap shows number of cell types in which each gene is dysregulated in. Genes are grouped and clustered based on hierarchical clustering and dendrogram clustering, respectively.

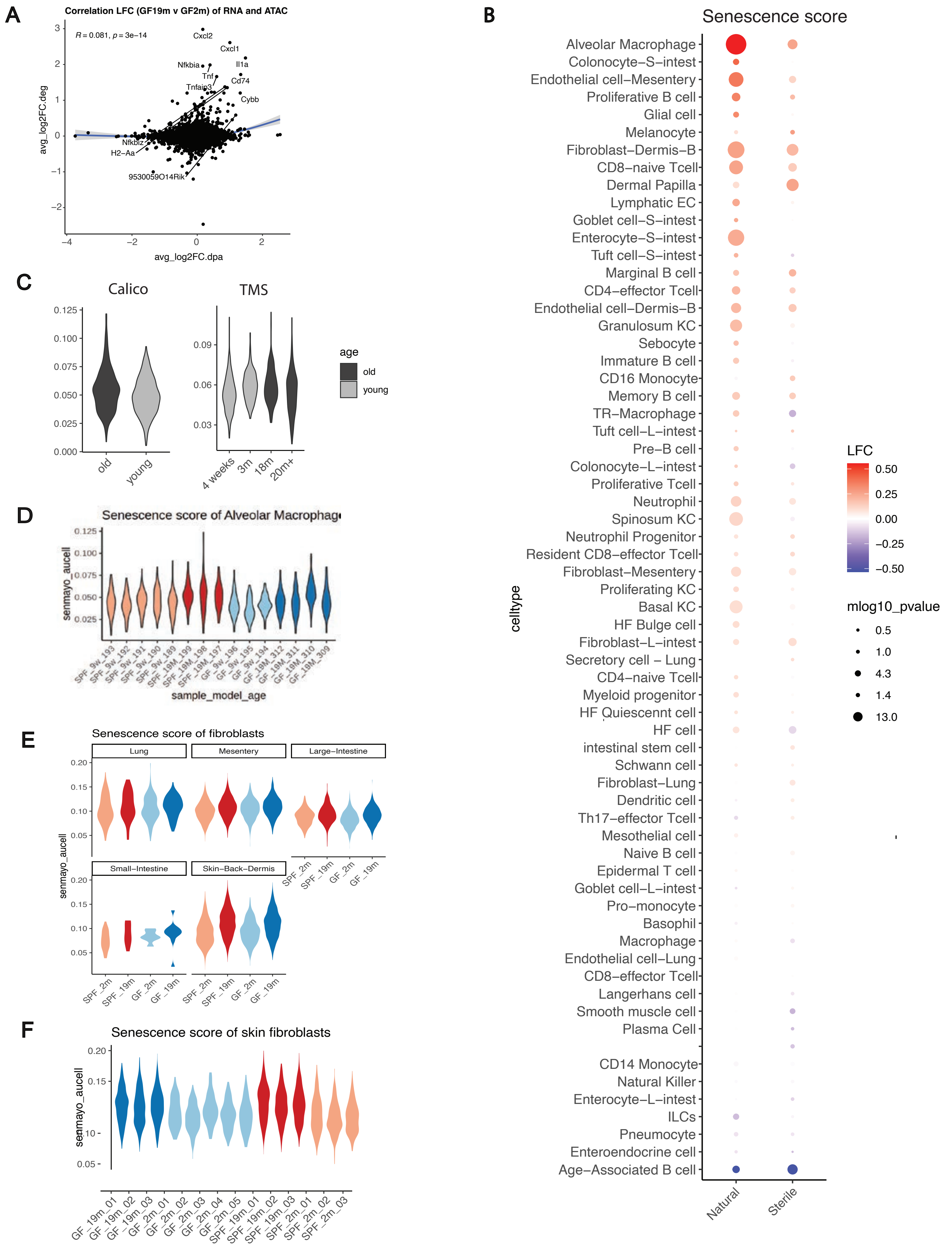


**Figure S4. Profiling of senescecent alveoloar macrophages. A.** Correlation of differential peaks to differential genes for alveolar macrophages in sterile ageing. For each peak, a gene was chosen based on the most accessible promoter peak. Scores are shown as the log2 fold change (LFC). **B.** LFC of senescence scores (mean score of Senmayo genesets, calculated with AUCell) in natural and sterile ageing across cell types. Significance is tested with Wilcoxon rank-sum test (P value <0.05). **C.** Senmayo scores of alveolar macrophages in the Tabula muris senis (TMS) and Calico ageing lung datasetss. **D-F.** Senescence score calculated with AUCell with Senmayo genesets of alveolar macrophages per mouse (D), fibroblasts across tissues (E) and dermal fibroblasts from skin per mouse (F).

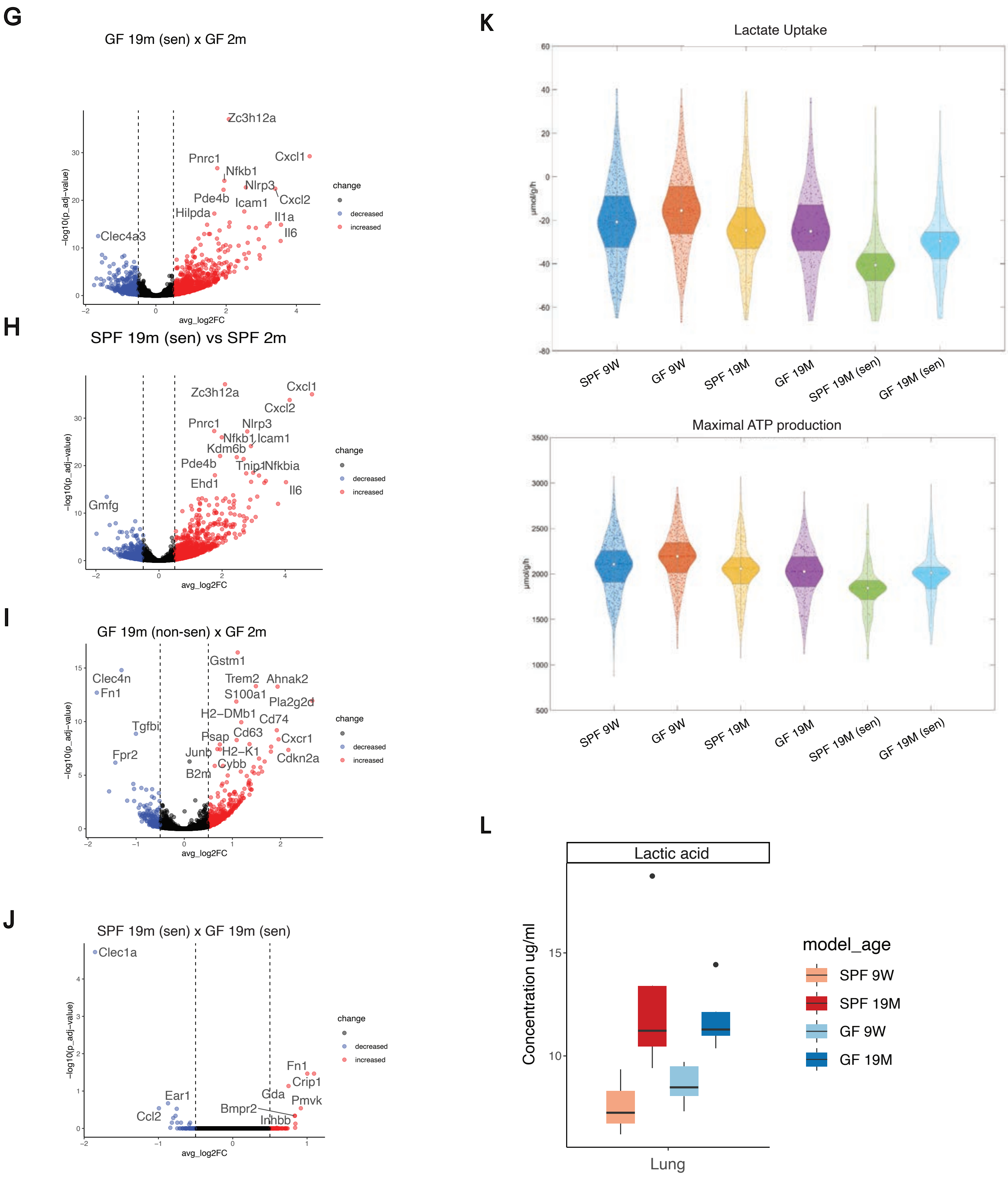


**Figure S4. Profiling of senescecent alveoloar macrophages (continued). G-J.** Volcano plots of DEGs between indicated subpopulations of alveolar macrophages. **K.** Violin plots depict the distribution of predicted metabolic functions derived from scRNA-seq data, showing a decline in lactake uptake and ATP production functions specifically in senescent alveoplar macropahge population. Shaded area indicates 25% and 75%. The dots indicate the metabolic functions for individual cells. Differences between groups are assessed by two-sided t-test for normal distribution or Wilcoxon signed-ranked test otherwise . Significance level Is indicated above brackets. **L.** Lactic acid levels from lungs measured by mass spectrometry shows an increase/ with old age in both natural and sterile ageing.

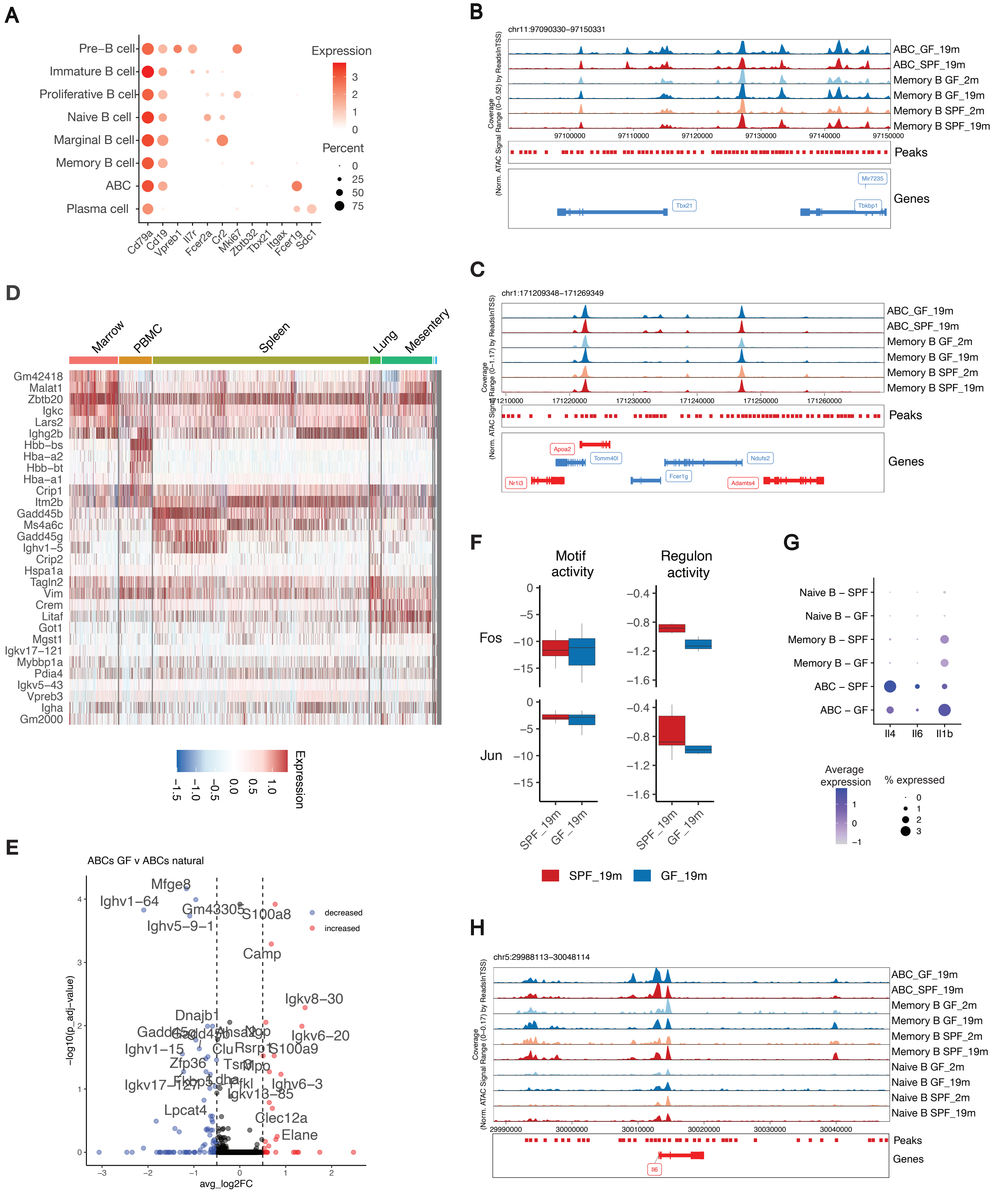


**Figure. S5A-H** (A) Marker genes for B cell subpopulations are visualised by their average expression per cell type. (B-C) Genome browser shot of Tbx21 (B) and Fcer1g (C) for ABCs (excluded young due to low count) and memory B cells. Reads were normalised by reads in TSS. (D) Gene expression heatmap of ABC genes that are differentially expressed between tissues. (E) DEGs between natural and sterile ageing of ABCs visualised by a volcano plot. (F) TF activities (motif on the left, regulon on the right) for Fos and Jun in ABCs. Boxplot represents score of all single cells. (G) Expression Dotplot of B cell subpopulations separated by condition and genes of interest. (H) Genome browser shot of Il6 locus for shown cell types. ABC_2m are omitted due to low cell count.Reads were normalised by reads in TSS.

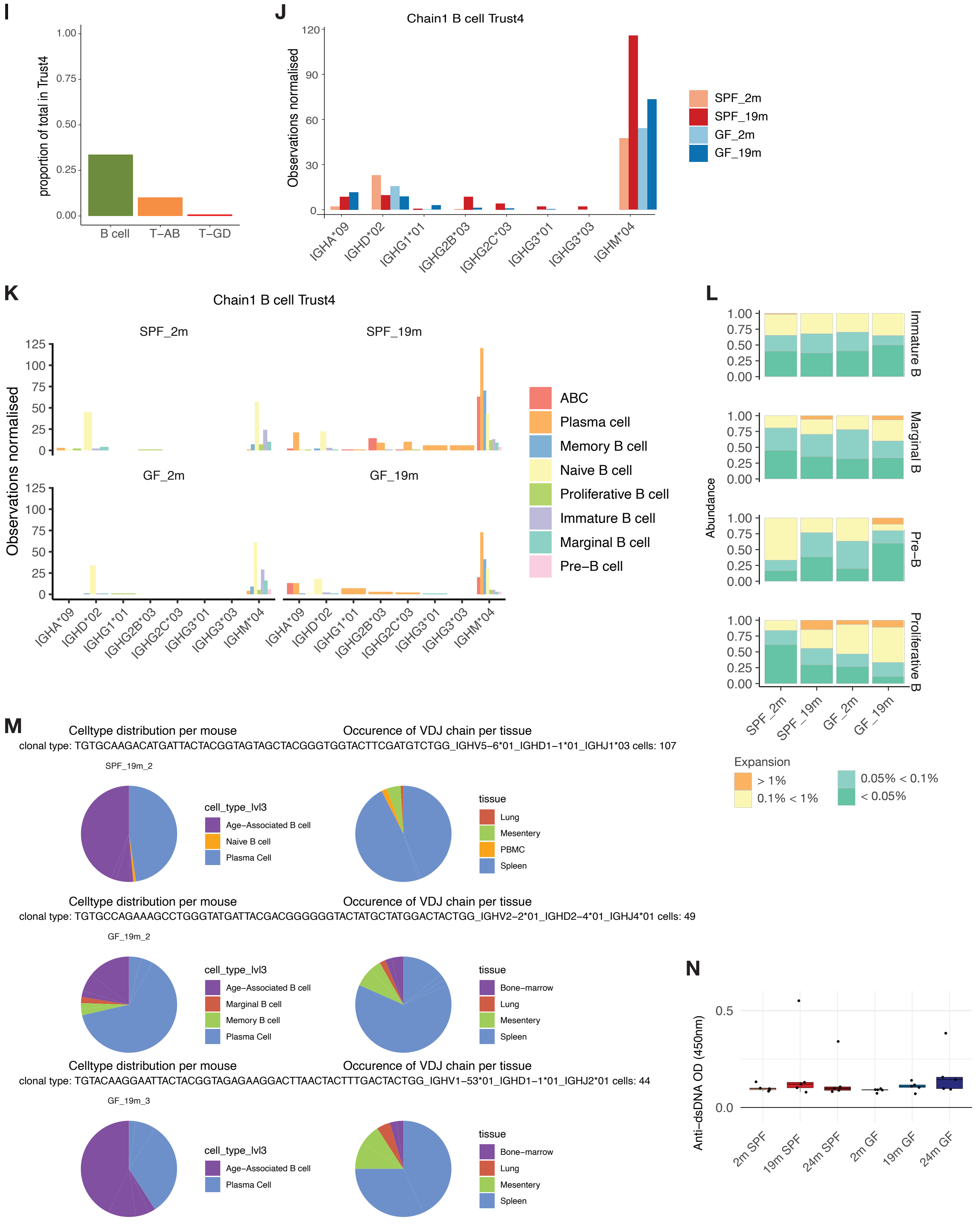


**Figure. S5I (continued).** (I) Observed lymphocytes in trust4 data. (J-K) B cell isotype annotations across conditions (J) and by B cell subpopulations (K). (L) Clonality analysis showing frequency of similar clones for B cell subpopulations. Occurrence is calculated per clonal type; degree of expansion is depicted by binning into >1% (orange), 0.1% < 1% (light green), 0.05% < 0.01% (green), and < 0.05% (dark green). (M) Example distributions of clonal typing across B cell subpopulations and tissues. (N) Auto-antibody levels of dsDNA in shown conditions measured by an ELISA assay. n=5
